# Perturb-ME: Scalable mechanism discovery from phenotype-enriched genome-wide screens

**DOI:** 10.64898/2026.08.17.745330

**Authors:** Hanchen Wang, Jiacheng Gu, Chris J. Frangieh, Michael S. Cuoco, Maryann Zhao, Arindam Sett, Tom Beyer, Kuan Pang, Etzard Stolte, Zihan Xu, Jure Leskovec, Orit Rozenblatt-Rosen, Vivek Natarajan, Kathryn Geiger-Schuller, Pratiksha I. Thakore, Aviv Regev

## Abstract

Perturb-seq enables pooled genetic screens with rich single-cell profiling readouts, but genome-scale profiling remains costly and may not be associated with other established functional characteristics. Moreover, as screens grow in size and complexity, interpreting the resulting data comprehensively is challenging and slow. Here, we introduce Perturb-seq with Marker Enrichment (Perturb-ME), which combines genome-scale CRISPR screening, phenotype-based enrichment and multimodal single-cell profiling. Applied to MHC-I cell surface protein expression in melanoma, Perturb-ME profiled HLA-low and HLA-high cells with matched RNA, surface-protein and guide measurements. A regulatory model with 221 impactful regulators affecting 1,998 responsive genes recovered seven coherent co-functional regulatory modules governing nine gene programs, including the canonical IFNγ-MHC-I axis regulating an antigen-presentation and interferon-response program. Agentic interpretation of the entire model with an AI co-scientist linked additional modules to trafficking, proteostasis and chromatin regulation. Perturb-ME, along with agentic interpretation, provide a scalable framework for comprehensive functional discovery from phenotype-enriched genetic screens.

## MAIN

Systematic mapping of causal molecular circuits is critical to understanding how disease phenotypes unfold and for identifying potential avenues for intervention. While pooled genetics screens are a powerful approach for such mapping, they were originally limited to low dimensional readouts, such as cell viability or the expression of a specific reporter protein, and thus could not assess the broader effects of each perturbation, nor could they relate perturbations effectively to each other, except as equivalent ‘hits’^1^. To address these limitations, Perturb-seq links genetic perturbations to high-content single-cell molecular profiles, such as single cell RNA-seq (scRNA-seq) enabling the simultaneous construction of genome-scale regulatory models in cells, organoids or organs *in vivo*, and opening the way to the construction of generalizable causal models of cells^1–4^.

However, applying Perturb-seq at genome and larger scales remains labor-intensive and cost-prohibitive experimentally and interpretation of the resulting massive datasets typically remains partial, despite intense efforts from human analysts. In a focused biological setting, only a subset of genome-wide perturbations is expected to alter the phenotype of interest, yet conventional genome-scale Perturb-seq distributes single-cell profiling uniformly across the full perturbation space. Moreover, most screens rely only on a cell’s RNA profile, but biological functionality is often associated with post-transcriptional phenotypes, such as the expression and localization of specific proteins. As a result, a large fraction of the experimental reagents and effort are spent on perturbations that are less informative for the phenotype of interest^5,6^. Existing approaches can reduce this burden by first performing a classical pooled enrichment screen and then constructing a focused library for a secondary single-cell screen, but this two-step strategy requires additional cloning, transduction and cell-culture cycles, slows down work considerably, and does not directly relate the molecular profile to the enrichment phenotype at the single cell level^7,8^. In addition, irrespective of the experimental strategies, even modest Perturb-Seq screens generate a treasure-trove of functional relationships in the form of the impact of each perturbed regulator on the expression level of each gene, which far exceed the assessment capacity of a human scientist, even for a single targeted screen. Although substantial efforts have been made to simplify this by analysis of global profiles^5,6^ or by decomposing perturbation effects into co-functional modules impacting gene programs^2,8^, none of these addresses the full nuance of the screen and converting these genetic effects into mechanistic hypotheses still relies heavily on manual and incomplete literature review and expert curation, and remains difficult to scale and prone to familiarity bias, where the most unusual and novel results may be ignored in favor of the previously established patterns.

Here, we introduce Perturb-seq with Marker Enrichment (Perturb-ME), an integrated strategy that combines genome-wide pooled CRISPR screening, marker-based phenotypic enrichment, and multimodal single-cell profiling through Perturb-CITE-Seq^9^ (**Fig. 1a**), and further combine it with an AI co-scientist for effective mechanistic interpretation. Perturb-ME enriches cells with altered surface expression of a phenotype-linked marker using fluorescence-activating cell sorting (FACS) and profiles these enriched cells by matched single-cell RNA, surface-protein, and guide-RNA sequencing. This design converts a genome-wide enrichment screen from a hit-discovery assay into a multimodal single-cell readout for mechanism discovery. To support interpretation, we further integrated Google DeepMind’s Co-Scientist^10^ into the analysis workflow. After quantifying perturbation effects and grouping perturbations and molecular features into regulatory modules and gene programs, respectively^2,9^, we provided the Co-Scientist with co-functional modules, gene programs, and enrichment-screen hits. The agentic system cross-referenced these results with existing literature to propose mechanistic annotations and prioritize hypotheses for follow-up, complementing statistical analysis by organizing high-dimensional perturbation effects into interpretable, testable biological hypotheses.

**Figure 1.**
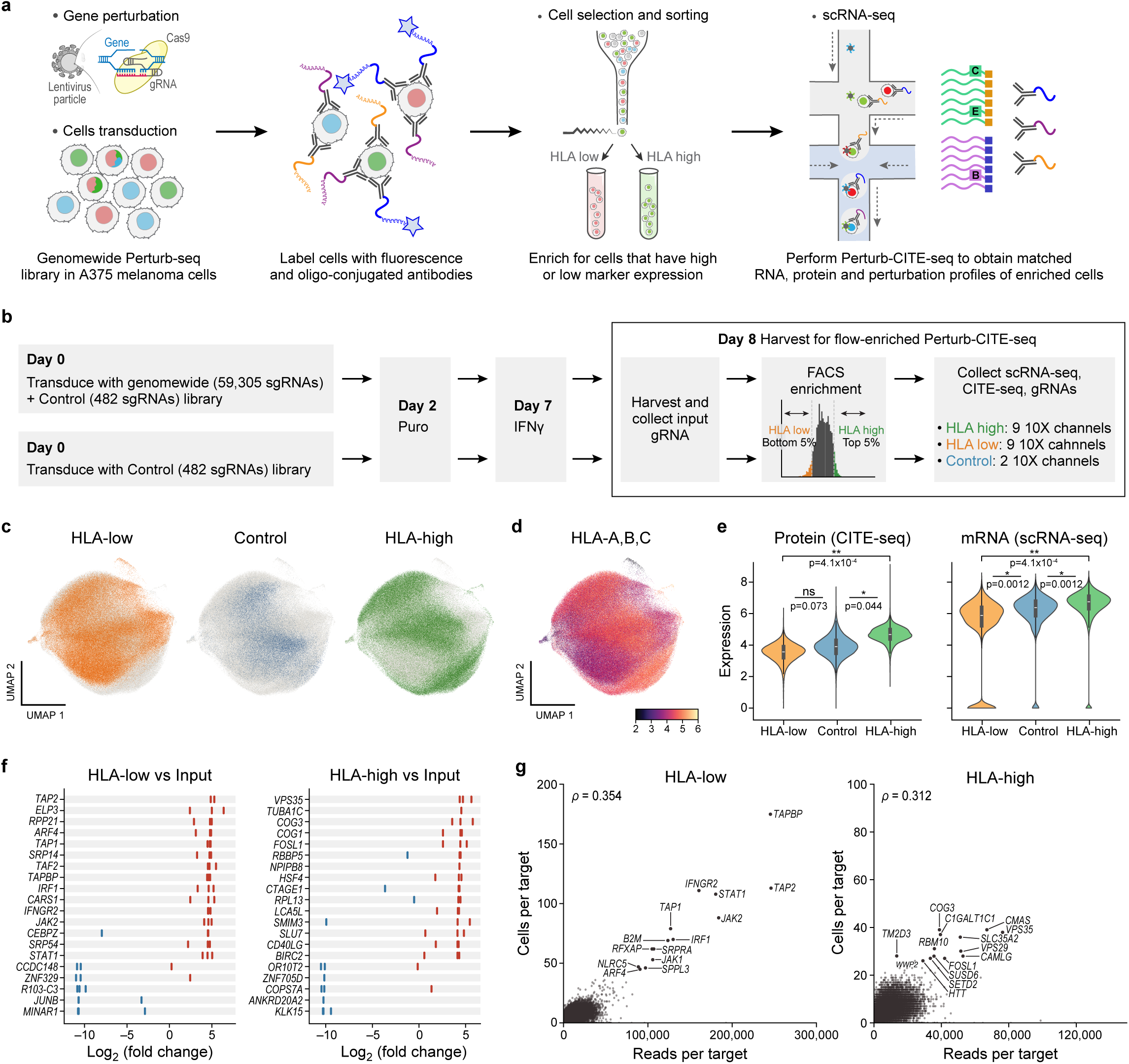
Perturb-ME enables genome-scale single-cell CRISPR screening of MHC-I regulators in melanoma. **a**, Experimental method overview. Cells are transduced with a genome-scale CRISPR-Cas9 knockout library (far left), stained with fluorescent anti-HLA-A/B/C antibodies to quantify HLA-I surface levels for FACS enrichment and with oligo-conjugated antibodies to measure target surface proteins by CITE-seq (second from left). HLA-low and HLA-high populations are sorted by flow (second from right) and profiled by Perturb-CITE-seq to recover matched single-cell RNA, surface-protein and sgRNA profiles (right). **b,** Perturb-ME screen in A375 melanoma cells. On day 0, A375 cells are transduced in parallel with the genome-scale library containing 59,305 sgRNAs (top left) or a control library containing 482 sgRNAs (bottom left). Cells undergo puromycin selection on day 2 (second from left) and IFNγ stimulation on day 7 (second from right), and are harvested on day 8 for input gDNA collection, FACS enrichment and Perturb-CITE-seq (right, box). **c-g,** Perturb-ME enriches phenotypically impacted cells. **c,d** UMAP embedding of single cell profiles (dots) colored by sorted population (c). or normalized surface HLA-A/B/C protein expression measured by CITE-seq (d). **e,** Distribution of HLA-A/B/C surface protein expression measured by CITE-seq (left, y axis) or HLA-A/B/C mRNA expression measured by scRNA-seq (right, y axis) in cell profiles from each bin (x axis). White dot, median; box, interquartile range.**f)** Change in gDNA *vs*. input gDNA library (x axis) for guides (tick marks) against each of the top enriched (red) or depleted (blue) targeted genes (rows, rank ordered) in HLA-low (left) or HLA-high (right) sorted populations. **g,** Number of single cell profiles assigned to each target in Perturb-ME (y axis) and number of bulk sgRNA reads per target (x axis) in HLA-low (left) and HLA-high (right) cells. ρ, Spearman correlation. Labeled points: selected top target perturbations from **f)**.

We applied Perturb-ME to identify regulators of MHC-I expression in the A375 melanoma cell line, representing a tumor type where loss of MHC-I (HLA-A/B/C) presentation on the malignant cell surface to CD8 T cells is causally associated with resistance to checkpoint immunotherapy^11^ (**Fig. 1a**). We chose A375 melanoma cells as a tractable IFNγ-responsive model and introduced either a genome-wide CRISPR knockout library of 59,545 guides targeting 19,769 genes (3 guides/gene) along with 482 control guides or, separately, a control-guide-only library, both at a low (<0.2) multiplicity of infection (**Fig. 1a,b**). We sorted cells carrying the genome-scale library into the bottom 5% and top 5% of HLA-A/B/C surface expression bins, corresponding to HLA-low and HLA-high populations, and collected cells carrying the control-guide-only library across the full HLA-I expression distribution to provide a matched control population (**Extended Data Fig. 3**). We profiled 172,578 HLA-low cells, 142,972 HLA-high cells and 37,541 control cells by Perturb-CITE-seq^9^, which links CRISPR perturbations to matched single-cell RNA-Seq and cell surface measurements of 13 protein markers through DNA-barcoded antibodies. The markers spanned canonical antigen presentation, immune regulation and cell-state proteins, providing protein-level phenotypes that complement mRNA abundance and are highly functionally interpretable (**Supplementary Table 1**). We recovered guide identities from expressed CROP-seq^12^ guide transcripts and assigned them to cell profiles through cell barcodes (**Methods**).

Phenotype-based enrichment substantially increased the effective single-cell coverage of perturbations impacting HLA-I surface expression. At the observed profiling depth of 277,152 cells, uniform sampling of the approximately 60,000-guide library in a conventional genome-wide Perturb-seq experiment is expected to yield approximately four sgRNA-singlet cells per guide, far below the typical number needed to assess robust effects^2,13,14^. In contrast, sgRNAs enriched in the HLA-low or HLA-high gates were recovered in tens to hundreds of cells, with the strongest hits represented by around 100 cells, whereas other sgRNAs were depleted (**Extended Data Fig. 5**). Perturb-ME therefore achieved up to 20-fold greater coverage of phenotype-associated perturbations, providing sufficient power to estimate their effects on RNA profiles and surface-protein expression.

Comparing HLA-low, HLA-high and control populations at both the RNA and protein levels showed that marker-based enrichment captured the intended HLA-I phenotypic states (**Fig. 1c-e**). In the joint embedding of single-cell RNA and protein profiles, HLA-low, control and HLA-high cells separated along an HLA-I expression axis (**Fig. 1c,d**). HLA-high cells had significantly higher HLA-A/B/C surface-protein and HLA-A mRNA expression than HLA-low cells (both P < 0.01, two-sided Mann-Whitney U test), with control cells intermediate (**Fig. 1e**). The CITE-seq cell surface protein panel revealed coordinated changes in additional proteins reflecting immune-regulatory and cell-state functions, including HLA-DR, CD274 (PD-L1), CD47, and CD49f, which were higher on the cell surface of HLA-high than HLA-low cells (P = 2.1 × 10^-3^, 6.7 × 10^-3^, 0.010 and 0.027, respectively; two-sided Mann-Whitney U test), whereas CD58 was unchanged (P=0.79) (**Extended Data Fig. 1**). This distinct behavior of CD58 is consistent with our previous study in patient derived melanoma cells, showing that CD58 is controlled by (and controls) different pathways than many checkpoints^9,15^ (Extended **Data Fig. 1**). These results confirm that phenotype-based enrichment recovered cells with the expected HLA-I states, while preserving matched RNA and cell surface protein profiles.

To compare Perturb-ME with a conventional bulk enrichment screen for cell surface expression of MHC-I, we measured the representation of bulk sgRNA in HLA-low and HLA-high populations relative to an input population collected before sorting and recovered 41 significantly enriched regulators (40 in the HLA-low and 1 in the HLA-high population)(**Fig. 1f**, FDR<0.05). These included known positive and negative regulators of MHC-I surface expression. For example, perturbations targeting IFNγ signaling components, including STAT1, IFNGR2, JAK2 and IRF1, were enriched as expected in the HLA-low population relative to input, consistent with the role of IFNγ signaling in transcriptionally activating MHC-I antigen-presentation genes^16^. Top HLA-low perturbations also included TAP1, TAP2 and TAPBP, core components of the MHC-I antigen-processing machinery required for peptide transport, peptide loading and stabilization of HLA-I complexes at the cell surface^17,18^. Conversely, the HLA-high population was enriched for perturbations of candidate negative regulators of HLA-I surface abundance, including VPS29 and VPS35, components of the retromer complex involved in endosomal sorting and recycling^19^. These trafficking-associated hits should not be interpreted simply as cargo-delivery factors; rather, they may reflect pathways that tune the balance between HLA-I transport, recycling and degradation, such that their disruption increases steady-state surface HLA-I levels. Consistent with this interpretation, prior genome-wide screens in HAP1 fibroblast-like cells and B cell lymphoma models have implicated antigen-processing, trafficking and endolysosomal pathways in MHC-I/II regulation^20,21^. Other less well-characterized candidates included conserved oligomeric Golgi complex genes COG1 and COG3.

Importantly, our phenotype-enriched single-cell profiles from Perturb-ME preserved the enrichment information measured by gDNA sequencing from the bulk pooled screens for both HLA-high and HLA-low phenotypes (**Fig. 1g**). In a conventional CRISPR guide enrichment screen, sgRNA read counts from amplified genomic DNA provide a proxy for the abundance of cells carrying each perturbation. In Perturb-ME, each profiled cell is directly linked to its perturbation, allowing target-level cell counts to be compared with bulk sgRNA abundance in the same sorted population. In both HLA-low and HLA-high populations, the number of single cells assigned to each perturbation were positively correlated with bulk sgRNA enrichment across all genes (Spearman’s rank correlation ρ = 0.31 and 0.35 for HLA-low and HLA-high, respectively, **Fig. 1g, Extended Data Fig. 2a**), and much more strongly for significant hits, **Extended Data Fig. 4**). For each perturbed gene (its sgRNAs pooled), we also compared, within each sorted bin, the bulk gDNA reads (entire bin), the number of single cells, and the total guide UMIs captured (**Extended Data Fig. 2**). The number of guide UMIs per cell is narrowly distributed (median ≈ 29, s.d. ≈ 7) and uncorrelated to the number of cells recovered (ρ = 0.03 and 0.04 in the two HLA gates, **Extended Data Fig. 2b**), suggesting that recovery is not driven by the amount of guide expressed or captured. Overall, the concordance indicates that Perturb-ME preserves the primary enrichment signal of a bulk CRISPR screen, while adding matched RNA and cell surface-protein profiles for phenotype-enriched perturbations in a highly efficient design.

### Linear modeling identifies regulatory modules controlling MHC class I expression

We next used the single-cell Perturb-ME profiles from the enriched high and low bins to quantify perturbation-specific molecular effects on MHC-I regulation (**Fig. 2a**). We assigned detected sgRNAs to cells, and restricted downstream analyses to the 277,152 cells with only one sgRNA species (singlets). Because Perturb-ME intentionally profiles HLA-low and HLA-high cells rather than uniformly sampling the full genome-scale library, target representation varied substantially across the two populations, with some perturbations strongly enriched in one phenotype-enriched population. For regulatory model fitting, we retained gene-level perturbations with at least 18 sgRNA-singlet cells in at least one of the two HLA gates (HLA-low or HLA-high; counted per gate; **Methods**). This yielded 221 gene-level perturbations spanning 663 guides (median 28 cells per perturbed gene, interquartile range 24-32; range 19-175 across all gates, **Methods**) and two pooled control categories (non-targeting and intergenic guides, pooled separately for the HLA-high and HLA-low gates; **Methods**). The overlap between this modeled set and the bulk screen was only partial. Of the 41 regulators significant in the bulk sgRNA screen (FDR < 0.05), 31 passed the single-cell coverage threshold and were modeled, whereas the remaining 10 fell below it in both HLA gates; these 10 “orphan” hits were not modeled but were revisited during mechanistic interpretation. Conversely, the other 190 of the 221 modeled perturbations were profiled by Perturb-ME but did not themselves reach bulk significance.

**Figure 2.**
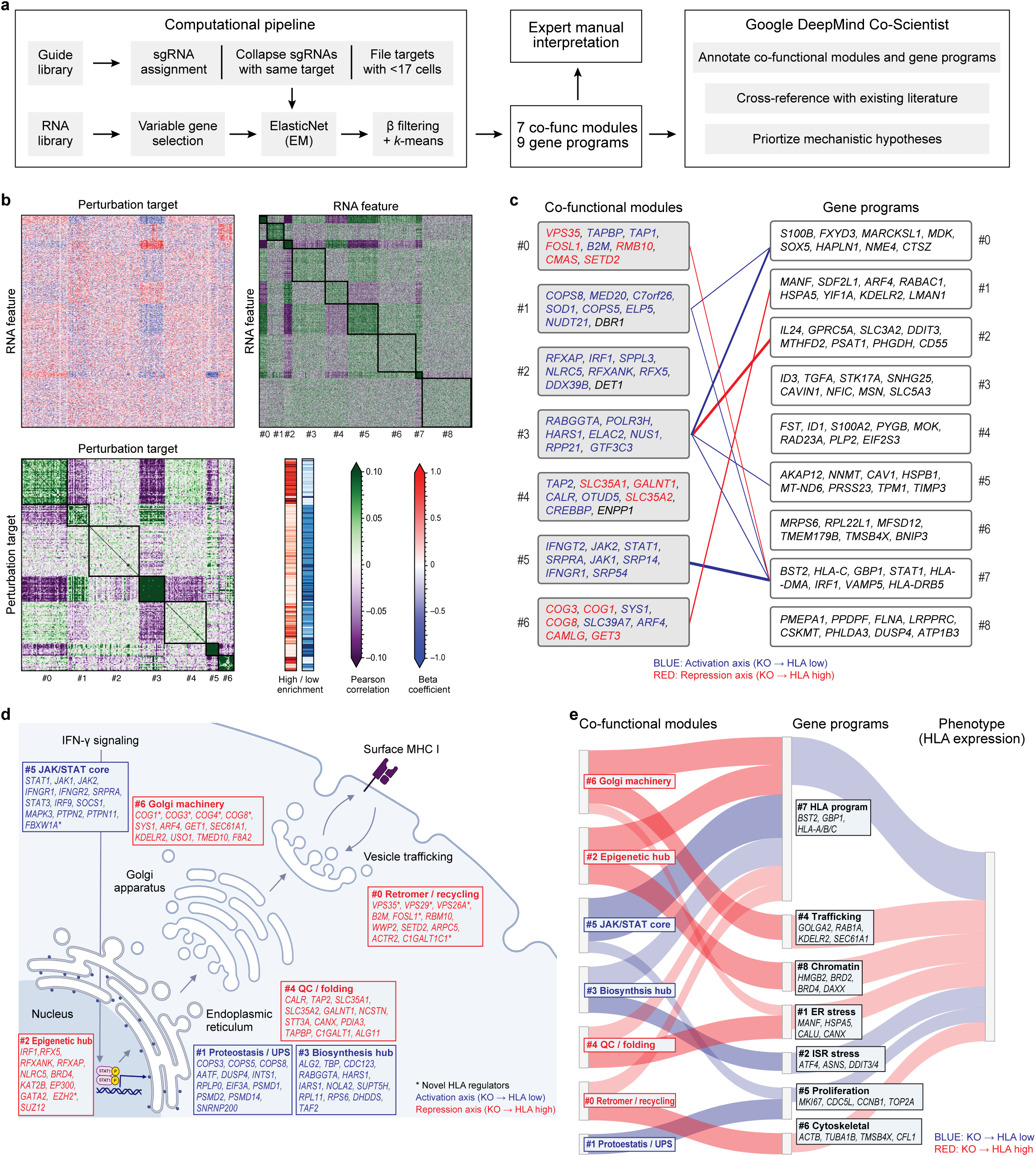
Linear-model decomposition and co-scientist interpretation of regulatory modules. **a**, Analysis workflow**. b,c,** Regulatory model of co-functional modules and co-regulated programs in the Perturb-ME screen. **b,** Top left heatmap: Regulatory effect size (β coefficient, red/blue color bar) for the impact on each RNA or protein feature (rows) of perturbing each gene (columns). Right and bottom matrices: Pearson correlation coefficient (purple/green color bar) between the significance profiles of either gene/protein features (right matrix) or perturbed genes (bottom matrix). Co-functional modules (bottom matrix) and co-regulated programs (right matrix) are identified by *K*-means clustering of each of the bottom and right matrices separately (*k*=7 and 9, respectively), and the clustering defines the row and column order. Narrow strips next to the target-correlation heatmap show single-cell counts per gene-target perturbation in HLA-High (red) and HLA-Low (blue), ordered identically to the target axis (**Methods**). **c,** Schematic of regulation of each program (right) by each module (left) and the regulatory logic (edge color; sign of mean β) and effect size (edge thickness, |mean β|). Edges with |mean β| < 0.15 are omitted (**Methods**). Eight representative genes shown for each program and module. Module genes are those with minimum MAGeCK pos-rank in either Low HLA or High HLA vs. Input. Program genes are those with the highest L2 norms of their β rows in the regulation matrix. Top-100 activators or repressors are colored blue and red, respectively. **d,** Co-Scientist interpretation of the co-functional modules across MHC-I biogenesis. Modules #0-#6 (boxes) mapped by Co-Scientist to candidate functional sites including IFNγ signaling, nuclear transcriptional regulation, ER biosynthesis and quality control, Golgi trafficking, retromer-dependent recycling and surface MHC-I expression. Representative members of each module are shown, colored by bulk MAGeCK enrichment for significant activators (blue), and repressors (red). Asterisks: candidate HLA regulators not previously implicated in the pathway. Edge color: activation (blue) or buffering/repression (red) axis (**Methods**). **e,** Co-Scientist-derived links between co-functional modules, gene programs and HLA-I surface protein. Proposed flow from co-functional modules, via gene programs to the HLA-High or HLA-Low phenotype. Flow color: sign of the mean β between each module and program (red/blue: up/down-regulation upon knockout). Flow width: aggregate effect magnitude (**Methods**).

To estimate direct perturbation effects while accounting for technical and biological covariates, we adapted the two-step regularized linear modeling framework used in prior Perturb-seq analyses^2,9^ to the Perturb-ME workflow (**Fig. 2a**; **Methods**). Briefly, we modeled log-normalized RNA and surface-protein features as a function of perturbation identity, cell-cycle phase, sequencing batch, sequencing depth and per-target coverage, with pooled control categories anchoring each sorted subpopulation (HLA-high, HLA-low). We estimated coefficients using ElasticNet followed by an expectation-maximization update that reweighted each cell’s perturbation assignment by its likelihood under the model. The response matrix related 221 gene-target perturbations and two pooled control categories to their effect on 18,683 molecular features (18,670 RNA transcripts and 13 CITE-seq surface-proteins). The model was based on 9,176 cell profiles and had 231 covariates (221 gene perturbations, two controls and eight technical or biological covariates) (**Methods**, **Supplementary File 4**).

We then used the regulatory submatrix of the effect of each gene perturbation on the mean expression of each RNA feature to identify seven co-functional regulatory modules across the 221 regulators with similar impacts across the RNA features, and nine programs across the 1,998 genes with similar RNA levels across the perturbations (**Fig. 2b, c**; **Methods**, **Supplementary Table 4**). This recovered a canonical IFNγ–MHC-I regulatory axis, where module #5, annotated as the JAK/STAT core, contained established regulators of IFNγ signaling and MHC-I induction, including STAT1, JAK1, JAK2, IFNGR1, IFNGR2, IRF9, SOCS1, MAPK3, PTPN2 and PTPN11. The model shows the expected result that perturbation of regulators in this module leads to down regulation of the HLA gene program (#7) which consists of antigen-presentation and interferon-response genes, including HLA-A/B/C, B2M, TAP1/2, TAPBP, GBP1, STAT1 and IRF1. This serves as a clear positive control of Perturb-ME’s ability to recover biologically coherent regulatory modules from phenotype-enriched single-cell profiles. Beyond this canonical axis, the same decomposition highlighted additional regulatory modules associated with biosynthesis, proteostasis, chromatin regulation, ER quality control, Golgi trafficking and retromer-dependent recycling.

### Agentic interpretation links Perturb-ME modules to candidate mechanisms

We next asked whether an agentic AI system could help convert Perturb-ME outputs into mechanistic hypotheses. We provided Google DeepMind’s Co-Scientist^10^ with three complementary data views: the seven co-functional modules from the regulation matrix, a stratified selection of the top 25 representative genes from each of the nine feature programs (to mitigate representation bias arising from skewed program sizes ranging from 61 to 461 genes), and an expanded, rank order list of the top 100 activator and top 100 repressor hits from the genome-wide bulk enrichment screen (by MAGeCK analysis^22^, **Methods**), to provide a broader context than the 41 significant ones from the bulk screen, which were all activators. We asked Co-Scientist to annotate the modules, cross-reference candidate regulators with existing literature, and propose links between co-functional modules, gene programs and the HLA surface-expression phenotype (**Supplementary File 2**). By cross-referencing the magnitude of phenotypic impact from the bulk screen with the causal expression signatures from single-cell profiles, the Co-scientist resolved the network into a bimodal “HLA Rheostat” mechanistic explanation (**Supplementary File 2**).

The Co-scientist framework evaluated competing mechanistic models through a tournament, ultimately synthesizing the winning hypothesis into a cohesive regulatory framework: the bimodal ‘Regulatory Rheostat,’ where a ‘stable epigenetic baseline volume’ is coupled with a ‘fast-acting post-translational degradation assembly line’ that dynamically tunes HLA-I surface density. The Co-scientist identified epigenetic control as a foundational ‘master switch,’ noting: ‘*Several hypotheses converge on the idea that epigenetic mechanisms are fundamental “master brakes” on the antigen presentation pathway… This direction suggests that the cell’s epigenetic state, often influenced by oncogenic signaling, is a primary determinant of its immunogenic potential*’ (Co-Scientist Summary, p. 2). It defined the system as a ‘*fragile*’ end-to-end assembly line, where ‘*disruptions in proteostasis, autophagy, or ER-Golgi transport represent critical points of failure that prevent functional HLA-I from reaching the cell surface, irrespective of transcriptional control*’ (p. 2). Finally, it resolved the observed data asymmetry (the extreme abundance (40:1) of activator hits compared to repressor hits in the bulk screen) by attributing active repression to oncogenic signaling, stating: ‘*The constitutive BRAF(V600E)-MAPK signaling pathway actively suppresses HLA-I expression… [through] rapid post-translational internalization and lysosomal sequestration*’ (p. 2). Thus, Co-Scientist interpreted this skewness as a biological outcome of tonic repression and deduced from the literature that because melanoma cells are already under chronic suppression (via constitutive BRAF signaling), knocking out a repressor yields a very weak rescue in HLA level, whereas knocking out any remaining activators causes a catastrophic collapse of HLA cell surface expression.

To develop this model, Co-Scientist first mapped the co-functional modules onto candidate cellular processes involved in MHC-I biogenesis (**Fig. 2d**). Modules of genes whose perturbation decreased HLA expression were organized into an activation axis spanning IFNγ/JAK-STAT signaling, protein synthesis and proteostasis. This axis included module #5, the canonical JAK/STAT core containing STAT1, JAK1, JAK2, IFNGR1, IFNGR2, IRF9, SOCS1, MAPK3, PTPN2, and PTPN11; module #3, a biosynthesis-associated module containing ALG2, TBP, CDC123, RABGGTA, HARS1, IARS1, RPL11, RPS6, DHDDS and TAF2; and module #1, a proteostasis/UPS-associated module containing COPS3, COPS5, COPS8, AATF, DUSP4, INTS1, EIF3A, PSMD1, PSMD2, and PSMD14. Together, these modules support a model in which IFNγ signaling, biosynthesis and proteostasis maintain antigen-presentation capacity.

Conversely, Co-Scientist organized modules with perturbations that increased HLA expression into a repression or buffering axis spanning chromatin regulation, ER quality control, Golgi trafficking and retromer-dependent recycling (**Fig. 2d**). This axis included co-functional module #2, an epigenetic hub containing RFX5, RFXANK, RFXAP, NLRC5, BRD4, KAT2B, EP300, GATA2, EZH2 and SUZ12; co-functional module #4, an ER quality-control and folding module containing CALR, TAP2, SLC35A1, SLC35A2, GALNT1, STT3A, CANX, PDIA3, TAPBP, C1GALT1 and ALG11; co-functional module #0, a retromer/recycling-associated module containing VPS35, VPS29, VPS26A, B2M, FOSL1, RBM10, WWP2, SETD2 and C1GALT1C1; and co-functional module #6, a Golgi-machinery module containing COG1, COG3, COG4, COG8, SYS1, ARF4, GET1, SEC61A1, KDELR2, USO1 and TMED10. These annotations suggest that HLA surface abundance is buffered not only by transcriptional regulation and antigen-processing pathways, but also by protein folding, trafficking and recycling processes.

Co-Scientist then linked these co-functional modules to downstream feature programs and the HLA surface-expression phenotype (**Fig. 2e, Methods**). In this analysis Co-Scientist connected the JAK/STAT module to an HLA/interferon-response program containing genes such as STAT1, IRF1, TAP1, TAPBP and HLA-A/B/C, while connecting Golgi, retromer and ER-associated modules to trafficking, ER-stress, ISR-stress, chromatin and cytoskeletal programs. In this representation, the modules provide perturbation-level regulatory units, the feature programs capture molecular response states, and the final phenotype summarizes whether perturbations shift cells toward HLA-low or HLA-high expression. Note that while we assembled the figure summarizing Co-Scientist’s findings (**Fig. 2e**), and the connections are the direct ones in the regulatory model (*i.e.*, were part of the input to Co-Scientist), Co-Scientist autonomously chose the modules-programs relationships to highlight and their interpretations individually and holistically.

Importantly, Co-Scientist’s interpretation not only recovered well-established biology but actively prioritized less-characterized mechanisms for follow-up (**Extended Data Table 1**). For example, Co-Scientist linked COG-complex and retromer components in modules #6 and #0 to altered HLA-I surface signal, highlighting undercharacterized, context-specific roles for Golgi and endosomal pathways in HLA-I surface homeostasis. It also highlighted ER quality-control factors in module #4, including CALR, CANX, PDIA3 and TAPBP, as candidate modulators of antigen-presentation machinery. Finally, the Co-Scientist prioritized chromatin-associated factors in module #2, including EZH2 and SUZ12, as candidate epigenetic regulators of HLA expression, consistent with prior work linking Polycomb repression to silencing of antigen-presentation genes^23^.

Beyond these module-level annotations, Co-Scientist generated specific literature-grounded hypotheses that connect individual hits to candidate regulatory routes. For example, it nominated FBXW1A in the JAK/STAT module as a potential repressor of the signaling core, proposed that perturbation of ER quality-control factors may expose stress-response branches that enhance antigen-presentation programs, and linked proteostasis-related regulators such as SQSTM1/p62 and NFE2L2 to stress-dependent modulation of HLA expression. As noted above, it also highlighted EZH2 as a candidate chromatin-level brake on MHC-I expression. These examples illustrate how the agent moved from hit lists and co-functional modules to more testable mechanistic models at the level of individual genes and pathways. The full set of Co-Scientist hypotheses is provided in the **Supplementary Material**.

We handle these Co-Scientist annotations as literature-grounded hypotheses rather than definitive mechanistic assignments. In this role, the agentic workflow complements the interpretable linear model: the statistical analysis estimates perturbation effects and identifies modules from direct experimental observations, whereas Co-Scientist helps organize those modules into candidate mechanisms that can be inspected, compared with manual interpretation and prioritized for experimental validation.

### Integrating statistical modules with agentic interpretation

Together, the linear model and Co-Scientist analysis define an efficient two-step workflow for functional screens with a high content readout: a focused, interpretable and efficient experimental workflow through phenotype-enriched perturbation profiles and a rapid, large-scale interpretation to yield mechanistic hypotheses for the consideration of the human scientist, which are grounded in the measured data and literature. The linear model provided a data-driven decomposition of perturbations and their effects, into modules and programs, while Co-Scientist organized these into literature-grounded, experimentally testable models of MHC-I regulation. A critical component of this workflow was expanding the Co-Scientist’s input to include a rank-ordered list of the top 200 MAGeCK hits (100 activators and 100 repressors), and not only the 41 significant (FDR<0.05) regulators from the bulk analysis, of which all but one were activators. This strategy provided the model with a more balanced regulatory landscape, enabling it to conceptually map onto the rheostat’s axes the 10 ‘orphan’ hits which were significant (FDR<0.05) in the bulk MAGeCK screen but excluded from the single-cell model due to the cell-coverage threshold, based on literature-derived mechanistic links. Had we only fed the Co-Scientist the 41 high-confidence hits (40 activators and 1 repressor), it would have been largely blind to the repressive axis. Expanding the list, along with the Perturb-ME regulatory model provided Co-Scientist with the ‘sub-threshold’ clues needed to spot the epigenetic baseline and derive the bimodal rheostat model.

Our study has several limitations. First, we studied a single melanoma model (A375) under cytokine-induced conditions, so the generality of the recovered HLA-I regulators to other tumor types and contexts remains to be tested. Second, marker-based enrichment concentrates statistical power on perturbations that alter the selected surface phenotype, and the linear model focused on the effect of single perturbations on the measured RNA and protein features rather than higher-order interactions or other phenotypic effects. Third, the reported hits depend on specific thresholds, including top-ranked perturbations in the bulk screen and a minimum of 18 single-cell singlets per perturbation for modeling, so weaker or sparsely sampled effects may be missed. Fourth, we only provided Co-Scientist with a subset of the program genes, to mitigate the imbalance in program size. Finally, the Co-Scientist produces literature-grounded hypotheses rather than direct evidence; its annotations are bound by the existing literature, may inherit its biases, and require experimental validation. Future work spanning additional cell types, perturbation modalities, and prospective tests of these hypotheses will help further establish the broader utility of Perturb-ME.

Together, these analyses highlight the complementary roles of statistical modeling and agentic interpretation in Perturb-ME. By combining phenotypic enrichment, multimodal single-cell profiling, perturbation-effect modeling and agentic interpretation, Perturb-ME provides a scalable framework for moving from genome-scale phenotypic screens to mechanistic hypotheses for further studies.

## EXTENDED DATA FIGURE LEGENDS

**Extended Data Figure 1.**
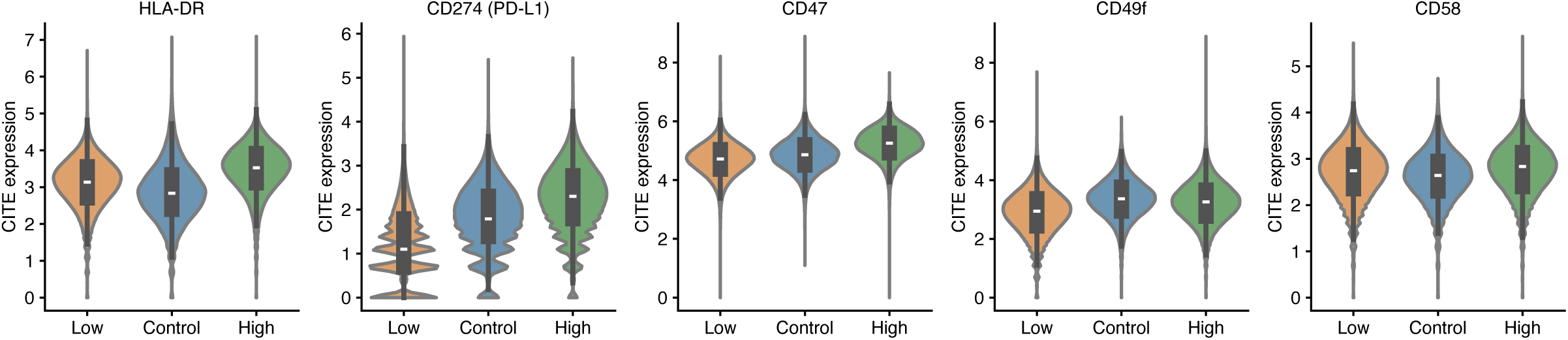
Increased surface-protein levels in HLA-high vs. HLA-low cells. Distribution of normalized surface-protein expression (y axis) from CITE-seq of different proteins (label on top) in HLA-low, control and HLA-high populations (x axis). White dot, median; box, interquartile range. *, Benjamini-Hochberg FDR < 0.05; **, FDR < 0.01; ns, not significant, two-sided Mann-Whitney U test on per-channel mean expression (*e.g.*, n = 9 HLA-high and 9 HLA-low 10x channels).

**Extended Data Figure 2.**
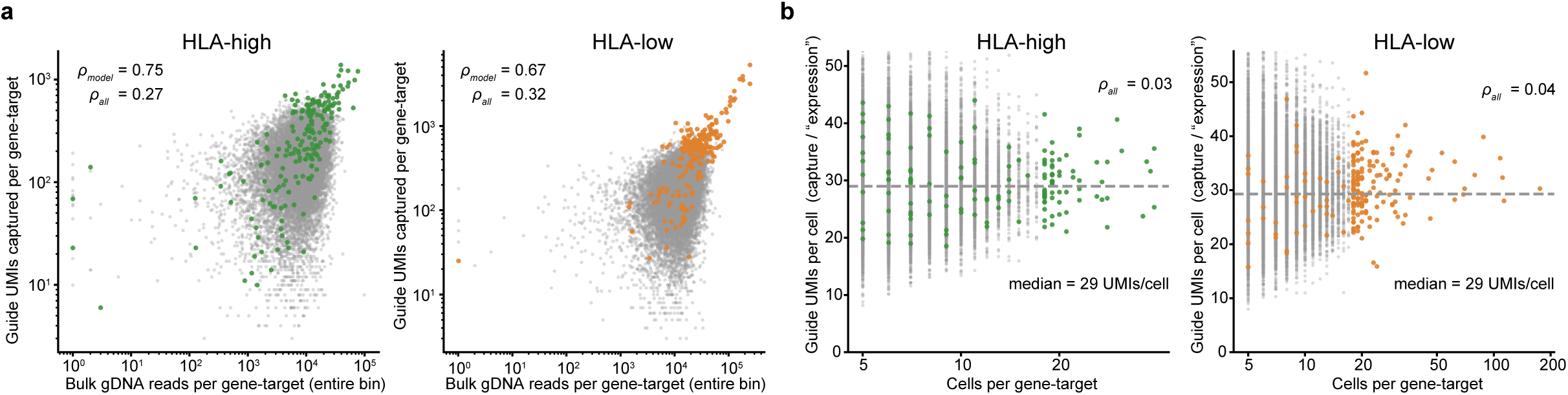
Comparison of perturbation detection by bulk gDNA reads, single-cell counts and captured guide UMIs within each HLA gate. **a,** Number of guide UMIs (y axis) and bulk gDNA reads (x axis) detected for each target (dots; aggregate across all guides) in the HLA-high (left) or HLA-low (right) bin. Colored dots: targets included in the regulatory model. Upper left corner: Spearman ⍴ for all targets or those in the regulatory model. **b**, Mean guide UMIs per cell (y axis) and number of cells (x axis) detected for each target (dots; aggregate across all guides) in the HLA-high (left) or HLA-low (right) bin. Colored dots: targets included in the regulatory model. Upper right corner: Spearman ⍴ for all targets or those in the regulatory model. Dashed line and upper right corner: median number of UMIs per cell.

**Extended Data Figure 3.**
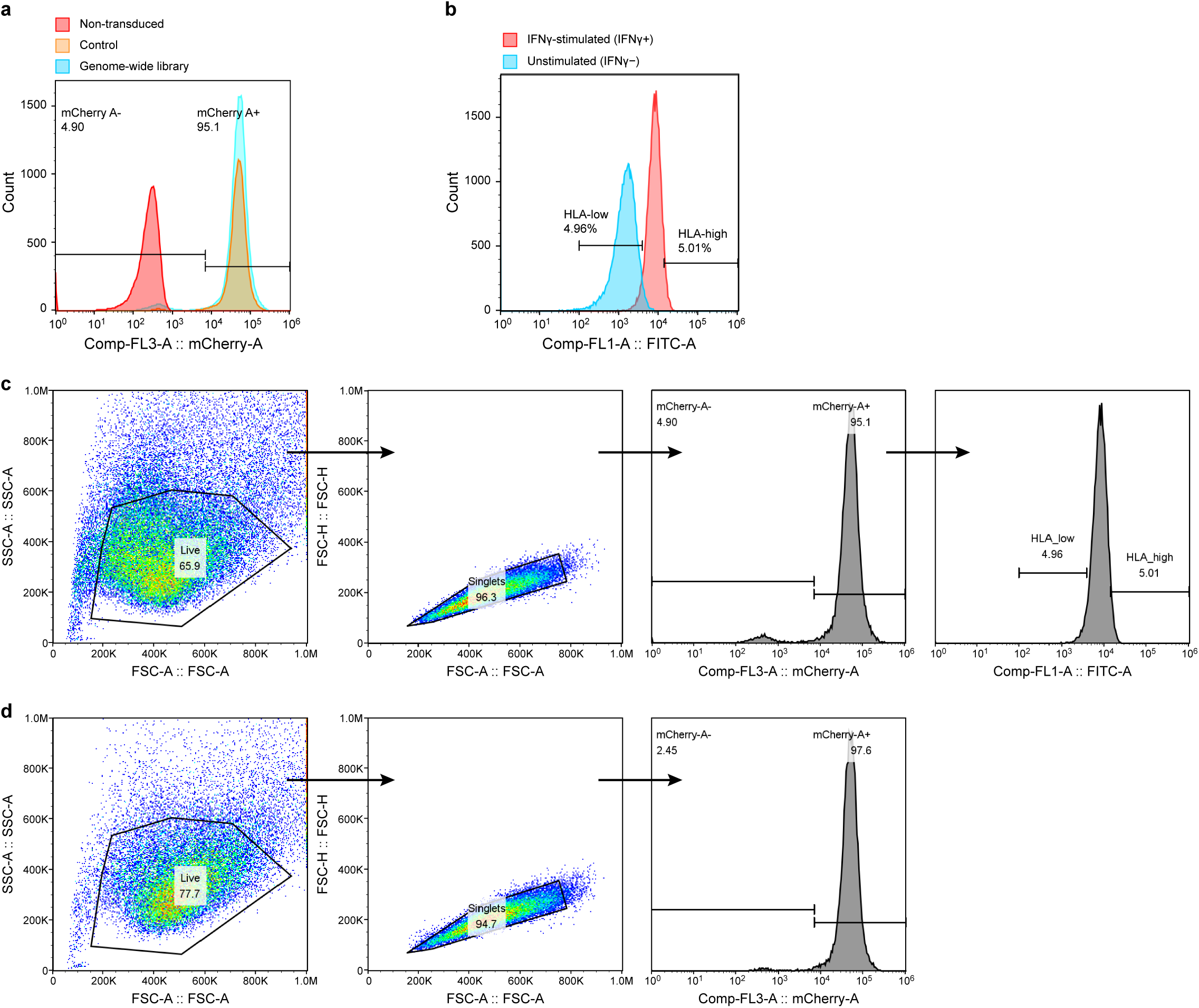
Flow cytometry gating strategy and enrichment sorting for Perturb-ME. **a**, Validation of CRISPR library transduction efficiency via mCherry marker expression. Distributions of mCherry fluorescence intensity in non-transduced cells (red) or cells transduced with the matched control library (orange) or the genome-wide library (cyan). In the genome-wide library sample, the mCherry-A− and mCherry-A+ fractions were 4.90% and 95.1%, respectively. **b**, HLA-A,B,C surface protein expression response upon IFNγ treatment. Distributions of FITC anti-HLA-A/B/C intensity (x axis) in IFNγ-stimulated cells (red) *vs*. unstimulated cells (blue). The HLA-low (4.96%) and HLA-high (5.01%) gates used for enrichment sorting are indicated. **c**, Sequential gating strategy for sorting genome-scale library samples (IFNγ-stimulated cells shown) into phenotype-enriched populations. Debris is excluded and intact cells are gated on FSC-A versus SSC-A (Live, 65.9%), followed by singlet discrimination on FSC-A versus FSC-H (Singlets, 94.0%). Transduced cells are isolated based on mCherry expression (mCherry-A+, 95.1%), and subsequently sorted on HLA-A/B/C (FITC) intensity to isolate the bottom ∼5% (HLA_low, 4.96%) and top ∼5% (HLA_high, 5.01%) populations for Perturb-CITE-seq. **d**, Sequential gating strategy for matched interferon stimulated, control-library samples (Ctrl-IFN2). Cells are gated sequentially for live intact cells (77.7%), singlets (94.7%), and mCherry-positive transduced cells (mCherry-A+, 97.6%) prior to profiling across the full unselected expression distribution.

**Extended Data Figure 4.**
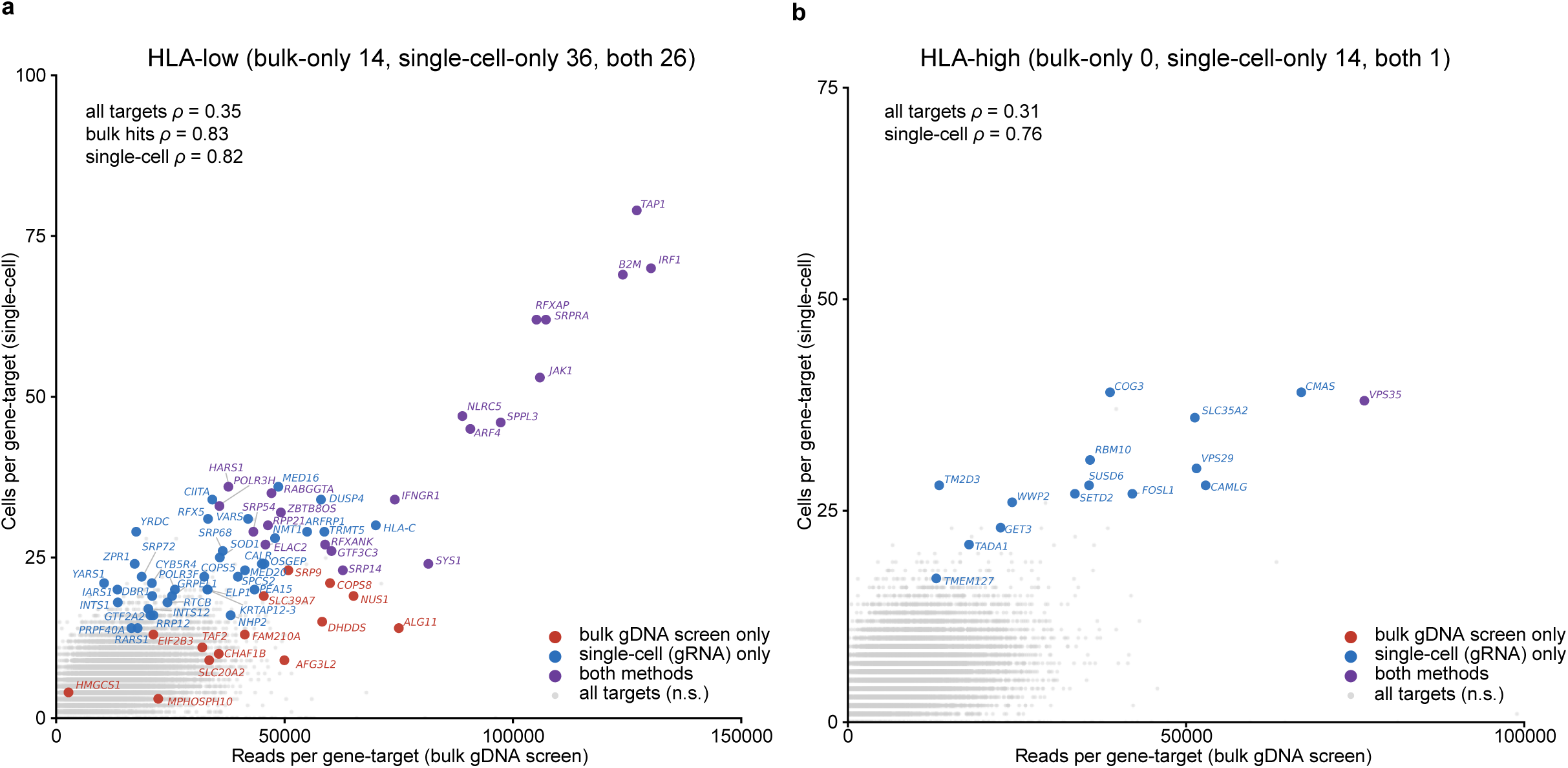
Agreement in significant hits between bulk gDNA screen and single-cell Perturb-ME enrichment. Number of reads per gene target in bulk gDNA flow-based bins (x axis) and number of single cells per gene target in Perturb-ME CITE-Seq (y axis), for the HLA-low (a) and HLA-high (b) analyses for each gene target (dot), called as a significant hit (FDR < 0.05) in bulk gDNA screen only (red; MAGeCK versus input), single-cell gRNA enrichment only (blue; genome-wide per-gene enrichment across the sorted gates, Methods), both (purple) or neither (grey). Upper left, Spearman ρ across all targets and among each method’s hits. Panel titles give the number of hits per category. Axes are cropped for clarity; five both-method HLA-low hits fall outside the plotted range (*TAPBP*, *TAP2*, *STAT1*, *JAK2*, *IFNGR2*; also labelled in Fig. 1g). Hit counts and Spearman ρ are computed over all hits, including those beyond the axes limits. n = 19,561 gene-targets.

**Extended Data Figure 5.**
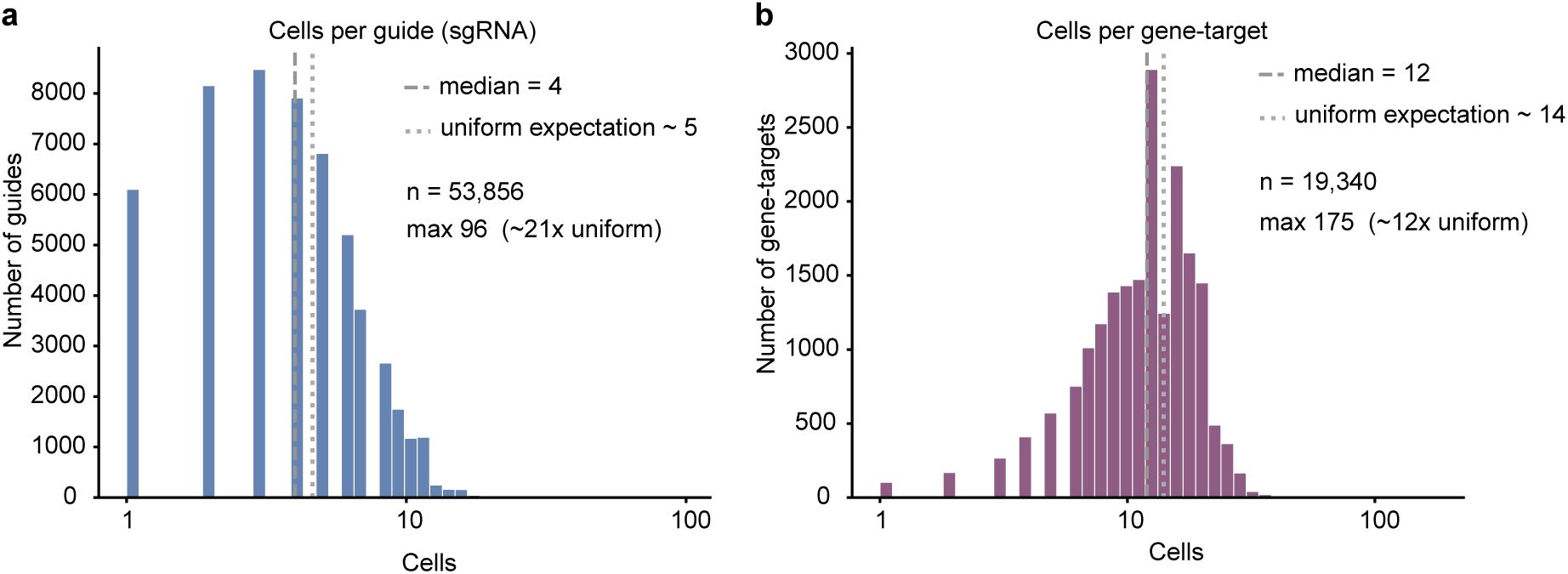
Number of single cells recovered per guide and per gene-target. Distribution of sgRNA-singlet cells recovered per guide (sgRNA) (a, x axis) or gene-target (b, x axis), across the ∼60,000-guide library (non-targeting and one-non-gene-site control guides excluded). Dashed line, median; dotted line, the number expected under uniform sampling of the library. Upper-left: total number and maximum (with its fold over the uniform expectation).

## METHODS

### Cell culture

A375 cells were obtained from the American Tissue Collection Center (ATCC) and were maintained in DMEM supplemented with 1% penicillin and 10% FBS. A Cas9-expressing A375 cell line was generated by treating A375s with lentivirus encoded with Cas9 and a blasticidin resistance gene. Lenti-Cas9-Blast. lentiCas9-Blast was a gift from Feng Zhang (Addgene viral prep # 52962-LV; http://n2t.net/addgene:52962 ; RRID:Addgene_52962). A375-Cas9 cells were selected and maintained in 6 ug/mL of Blasticidin.

### Generation of the pooled genome-wide Perturb-Seq library

A 59,545 single guide (sgRNA) library targeting 19,769 genes was designed, with 3 guides targeting each gene. Guide sequences were picked from the Broad Institute Genetic Perturbation Platform Web sgRNA Designer (https://portals.broadinstitute.org/gppx/crispick/public; **Supplementary Table 2**). An oligo array containing these sgRNA sequences as well as appropriate overhangs for Golden Gate cloning was synthesized (Twist Biosciences). A second library was created with 241 control non-targeting and 241 intergenic guides only, also through a synthetic oligo array (Twist Biosciences). The oligo arrays were amplified by PCR and cloned into a previously published backbone vector CROP-seq-mKate2 as previously described^9^. The genome-wide targeting gRNA library and control libraries were combined to generate the plasmid pool for the genome-wide Perturb-Seq experiment. The coverage and distribution of the genome-wide + control library and control library alone were assessed by PCR amplification and deep sequencing using an Illumina MiSeq.

### Lentivirus production and transduction

VSV-G pseudotyped lentivirus was generated in HEK293T cells cultured in high-glucose DMEM (Gibco, 11995) supplemented with 10% FBS and 1% penicillin-streptomycin. 18 h after seeding, cells in 10-cm plates were cotransfected with the gRNA library (12 μg), the second-generation packaging plasmid psPAX2 (Addgene, 12260, 10 μg), and the envelope plasmid pMD2.G (Addgene, 12259, 5 μg) using X-tremeGene 9 transfection reagent (Roche) according to manufacturer’s instructions. After 16 h, the transfection medium was exchanged for 10 mL of fresh 293T medium. Conditioned media containing lentivirus was collected 24 and 48 h after the first media exchange and combined. Lentivirus supernatant was filtered using a 0.45 μm filter and stored at -80°C prior to use. A375 cells were incubated with lentivirus supernatant for 24 hours in the presence of 4 μg/mL polybrene for transduction.

### Perturb-Seq with phenotype enrichment for HLA expression

A375 cells stably expressing Cas9 were transduced with lentivirus expressing the genome-wide + control gRNA library or the control gRNA library alone at a multiplicity of infection of 0.2, as determined by titering. Media was exchanged after an overnight incubation and transduced cells were selected with 2 μg/mL puromycin 72 h after initial treatment with lentivirus. 6 days after transduction, cells were treated with 2 ng/mL of recombinant IFNg for 24 h to induce HLA expression prior to harvest for flow enrichment prior to Perturb-CITE-Seq.

For harvest, cells were trypsinized and washed in PBS. Cells for bulk gRNA enrichment from genomic DNA were pelleted and snap-frozen. For Perturb-CITE-seq, cells were blocked in 2% FBS in PBS and incubated for 20 min on ice. To enable high-throughput overloading of droplet-based scRNA-seq with doublet detection, cells were also split into 6 sets and tagged with hashing antibodies against ubiquitous cell surface antigens. Cells were incubated for 30 min on ice in an antibody cocktail of fluorophore conjugated anti-human HLA-ABC (1:400, Biolegend), CITE-seq antibodies (**Supplementary Table 1**), and 1 of 6 barcoded hashtag antibodies (Biolegend, **Supplementary Table 1**). After incubation, cells were washed in a blocking buffer twice and the 6 populations were combined into a single pool prior to sorting. Cells were sorted on two Sony MA900s concurrently. Live cells were gated on an mKate+ population to select for cells expressing the gRNA library and then gated on HLA expression. For control library only-treated cells, the entire distribution of HLA expression was sorted. For the genome-wide + control library-treated cells, the lowest 5% and highest 5% cells based on HLA expression were sorted for Perturb-CITE-Seq and bulk sgDNA sequencing from genomic DNA.

For scRNA-seq, 40,000 cells were loaded onto each channel using the 10X Chromium system with the Chromium Single Cell 3’ Library and Gel Bead kit v.3.1 (10X Genomics) with the goal of recovering approximately 20,000 single cell expression profiles. 18 channels were loaded from cells treated with the genome-wide + control gRNA Perturb-seq library (9 channels from the bottom 5% HLA sorting gate (HLA low) and 9 channels from the top 5% HLA sorting gate (HLA high)), along with 2 channels from cells treated with the control gRNA library alone.

### Generation of the single-cell transcriptome, antibody, and guide RNA library

Single cell gene expression libraries were generated following the manufacturer’s instructions (10X Genomics) with modifications to generate the hashing, CITE-seq, and CROP-seq gRNA library. During cDNA amplification, 2 μL of the 2 pmol HTO additive primer and 3 μL of the 2 pmol of the ADT additive primer was spiked into the reaction to increase the abundance of hashtag antibody oligos and CITE-seq oligos. During purification of the cDNA amplification reaction, a 0.6X SPRI was performed to separate large gene expression library cDNA from smaller hashing and CITE-seq libraries. The supernatant of this 0.6X SPRI was set aside to generate the hashing and CITE-seq expression libraries. The 0.6X cDNA amplification SPRI was then completed and gene expression libraries were generated following the manufacturer’s instructions.

For the hashing and CITE-seq libraries, 1.4X SPRI beads were added to the initial 0.6X SPRI supernatant described above. This double sided SPRI was completed following two 200 µL 80% ethanol washes. The product was eluted in a 50 µL elution buffer (Qiagen). A final 2.0X SPRI was performed on this hashing and CITE-seq template with elution in 50 µL elution buffer (Qiagen). To generate the hashing sequencing libraries, 4 µL of the SPRI-cleaned template was mixed with 25 µL of 2X NEBNext HiFi Master Mix, 11.5 µL of diH2O, and 1 µL of a 1:1 HTO primer mixture (10 µM stock concentration of each primer). To enable multiplexing, one of three unique P7 indices were included in each reaction. The hashing sequencing library reaction was cycled with the following conditions using a thermocycler: 98°C for 10 sec, (98°C for 2 sec, 72°C for 15sec) x (22-26 cycles) followed by a final extension of 72°C for 1 minute. Products were purified by 2.0X SPRI clean-up and size of amplicons verified by Agilent Tapestation.

To generate the CITE-seq sequencing libraries, 5 µL of the SPRI-cleaned template was mixed with 25 µL of 2X NEBNext HiFi Master Mix, 11.5 µL of diH2O, and 1 µL of a 1:1 ADT primer mixture (10 µM stock concentration of each primer). To enable multiplexing, one of three unique P7 indices were included in each reaction. The CITE-seq library reaction was cycled with the following conditions using a thermocycler: 98°C for 10 sec, (98°C for 2 sec, 72°C for 15sec) x (19-24 cycles) followed by a final extension of 72°C for 1 minute. Products were purified by 2.0X SPRI clean-up and size of amplicons verified by Agilent Tapestation.

Amplification of fragments containing the guide barcodes was performed as described previously^9^. Briefly, a 10 ng fraction of the WTA was amplified using primers specific to the CROP-seq-mKate2 gRNA cassette. A second PCR was performed to attach Nextera adapters.

Libraries were sequenced on an Illumina HiSeq X to 15,000 reads/cell for scRNA-seq, 4,500 reads/cell for CITE-seq, and 1,000 reads/cell for hashing libraries. CROP-seq gRNA libraries were sequenced on an Illumina Nextseq 550 at approximately 2,000 reads/cell.

### Generation of bulk guide RNA library from genomic DNA

The bulk gRNA library was generated from genomic DNA as previously described^9^. Bulk gRNA libraries were generated from input cells collected prior to the sort, from cells sorted for lowest 5% of HLA expression, and from cells sorted for highest 5% of HLA expression. Genomic DNA was extracted and purified from cells using the QIAamp DNA Micro Kit (Qiagen, no. 56304) using no more than 500,000 cells per column. sgRNA sequences were amplified from genomic DNA for Illumina sequencing first using primers specific to the integrated portion of the CROP-seq-mKate2 gRNA expression cassette and then with primers designed to introduce Illumina sequencing compatible sequences. Libraries were sequenced on an Illumina HiSeq 2500 in RapidRun mode with custom read 1 (primer 503 F). All primer sequences are listed in **Supplementary Table 5**.

### Bulk guide RNA enrichment analysis

To quantify sgRNA enrichment in FACS-sorted populations, bulk guide RNA sequencing reads from genomic DNA were aligned to the sgRNA library and counted using MAGeCK v0.5.9.2^22^. Read counts were generated across nine samples: four input library replicates (ICR47_1A–D), two HLA-High replicates (High_A, High_B), two HLA-Low replicates (Low_A, Low_B) and one unsorted control (CTL). Across all samples, 58,802 sgRNAs were quantified with a median mapping rate of 91-95%. Counts were normalized using the median-ratio method implemented in MAGeCK.

MAGeCK tests were performed for three comparisons: HLA-Low versus Input, HLA-High versus Input and CTL versus Input, using the four input replicates as controls. sgRNAs with zero counts in both treatment and control groups were excluded, corresponding to 267 sgRNAs for the HLA-Low comparison and 278 sgRNAs for the HLA-High comparison. sgRNAs with extremely high counts, defined as more than four standard deviations above the mean, were flagged as outliers and excluded from variance estimation, corresponding to 146 and 150 sgRNAs for the HLA-Low and HLA-High comparisons, respectively. Gene-level enrichment and depletion significance were calculated using MAGeCK robust rank aggregation. sgRNA-level log2 fold changes were summarized at the gene level by taking the median across targeting sgRNAs.

Positive enrichment in the HLA-Low population indicates perturbations whose knockout decreases surface HLA-I abundance, whereas positive enrichment in the HLA-High population indicates perturbations whose knockout increases surface HLA-I abundance. HLA activators were defined as gene-target perturbations ranked in the top 100 by “pos|rank” in the HLA-Low versus Input comparison, and HLA repressors were defined as gene-target perturbations ranked in the top 100 by “pos|rank” in the HLA-High versus Input comparison. These bulk classifications were used to interpret HLA-I regulatory axes, to compare bulk guide enrichment with single-cell representation and to color gene labels in downstream regulatory-network visualizations.

For concordance analysis between bulk sgDNA enrichment and single-cell recovery, reads per gene were aggregated across sgRNAs targeting each gene and compared with the number of single cells assigned to the same gene target in the corresponding sorted population. Spearman correlation coefficients were computed separately for HLA-Low and HLA-High populations.

### CITE-Seq preprocessing and feature normalization

Raw single-cell sequencing data from 20 10X Genomics channels, including 9 HLA-Low channels, 9 HLA-High channels and 2 control channels, were processed with Cell Ranger (10x Genomics) to generate gene-expression and feature-barcode count matrices. Downstream preprocessing was performed using Cumulus^24^. Cells were demultiplexed using hashtag oligonucleotide barcodes, and only hashtag singlets were retained. Only cells with 800-6,000 detected genes and no more than 15% mitochondrial reads were retained. Genes detected in fewer than 100 cells across the dataset were removed, yielding 18,670 RNA features.

RNA counts were normalized to a target sum of 10^6^ counts per cell and log-transformed as log(x + 1). For protein measurements, raw antibody-derived tag counts were normalized against matched isotype-control antibodies as max(0, log((ADT count + 1) / (isotype-control count + 1)))^9^. Surface HLA-I protein abundance was quantified using normalized HLA-A/B/C CITE-seq signal, and HLA-I transcript abundance was quantified from the corresponding scRNA-seq features.

A shared UMAP embedding was generated from the preprocessed single-cell expression profiles and used to visualize HLA-Low, control and HLA-High populations. The same UMAP coordinates were used to display sorted population identity and normalized HLA-A/B/C protein abundance. Distributions of HLA-A/B/C surface protein and mRNA expression were compared across HLA-Low, control and HLA-High populations.

### sgRNA assignment and target-level perturbation representation

sgRNA identities were assigned to individual cells by aligning CROP-seq guide reads to the sgRNA reference, requiring at least 1 UMI for each sgRNA assignment. sgRNA assignment multiplicity was defined as the number of distinct perturbation targets detected in each cell. The final AnnData object contained 353,091 single-cell profiles passing RNA and hashtag-based singlet filters, with annotations for sorting condition, sample identity, sgRNA assignment, sgRNA assignment multiplicity, cell-cycle phase, sequencing run and 13 normalized CITE-seq protein measurements.

For perturbation-effect modeling, only sgRNA singlets were retained, defined as cells assigned with a single perturbation target. This yielded 277,152 sgRNA-singlet cells. sgRNAs targeting the same gene were collapsed to target-level perturbations. To focus on perturbations with sufficient single-cell profiles, only gene-target perturbations represented by at least 18 sgRNA-singlet cells in either the HLA-Low or HLA-High population were retained. This filtering step yielded 221 gene-target perturbations for downstream linear modeling.

### Linear modeling of perturbation effects

To quantify the molecular impact of individual perturbations, a regularized linear model was fit, relating perturbation identity to RNA and CITE-seq feature abundance, adapting the MIMOSCA approach^2^. The model was fit on sgRNA-singlet cells from the HLA-Low and HLA-High populations after perturbation-coverage filtering. This yielded 9,176 cells for model fitting, including 6,911 cells assigned to 221 gene-target perturbations and 2,265 cells assigned to pooled control categories. Non-targeting (“NO_SITE”) and intergenic (“ONE_NON-GENE_SITE”) control sgRNAs were pooled separately for each sorting condition to define “HIGH_CTL” and “LOW_CTL”.

The covariate matrix X contained 9,176 cells and 231 covariates. These covariates included binary indicators for 221 gene-target perturbations and two pooled control categories, together with cell-cycle phase, sequencing run, total UMI count and cells per target. The response matrix Y contained 18,683 features, comprising 18,670 log-normalized RNA features and 13 normalized CITE-seq surface-protein features.

An ElasticNet regression model was fitted independently for each feature, using the same *L*1 and *L*2 penalty ratios (0.5), a regularization strength of α = 0.0005, a precomputed Gram matrix and a maximum of 10,000 iterations. The resulting coefficient matrix β contained 18,683 features and 231 covariates. Positive β coefficients indicate that a perturbation increased feature abundance, whereas negative β coefficients indicate that a perturbation decreased feature abundance.

To refine sgRNA-to-perturbation assignments, an expectation-maximization-style update was applied. For each gene-target perturbation, a soft assignment probability was computed for each cell based on the improvement in residual sum of squares when the perturbation covariate was included versus excluded. The covariate matrix was then updated by replacing binary perturbation assignments with these posterior probabilities, and the ElasticNet model was refit. This EM refinement was applied only to the 221 gene-target perturbation covariates; pooled control and technical covariates were held fixed. The final EM-refined coefficient matrix was used for all downstream target-module, gene-program and regulatory-network analyses.

### Identification of co-functional modules and gene programs

To identify coordinated perturbation and response patterns, the EM-refined perturbation-effect matrix was clustered. Clustering was restricted to gene-target perturbations and RNA features with substantial regulatory effects. RNA rows in which fewer than 25% of coefficients had |β| ≤ 0.05 and gene-target columns in which fewer than 80% of coefficients had |β| ≤ 0.05 were retained, yielding a 1,998 × 221 perturbation-effect submatrix. CITE-seq surface-protein features were included in the ElasticNet response matrix but were not used to define RNA gene programs.

Pairwise Pearson correlation coefficients were computed across gene-target columns to obtain a target-similarity matrix and across RNA-feature rows to obtain a feature-similarity matrix. *K*-means clustering was applied independently to each correlation matrix using scikit-learn KMeans with n_init = 10 and random_state = 3. This defined seven co-functional modules and nine RNA gene programs.

The numbers of clusters were selected by joint inspection of correlation heatmap structure, within-cluster coherence and recovery of known HLA-I regulatory biology, including IFNγ/JAK-STAT signaling and MHC transcriptional regulation. A silhouette-score sweep did not identify a meaningful optimum, with scores near zero across tested *k* values, consistent with a continuous and gradient-like regulatory architecture rather than well-separated globular clusters. *k* = 7 co-functional modules and *k* = 9 gene programs were used to preserve interpretable biological structure while avoiding excessive fragmentation.

### Representative genes and module-program links

For the target-module and gene-program diagram, representative genes were selected separately for each side of the network. For each of the seven co-functional modules, members were ranked by the minimum “pos|rank” between the HLA-Low versus Input and HLA-High versus Input MAGeCK comparisons, with ties broken by the *L*2 norm of the corresponding β column in the EM-refined coefficient matrix. If a module contained fewer than eight members among the top 100 MAGeCK hits in either comparison, the list was completed using remaining module members with the largest β column *L*2 norm. For each of the nine gene programs, representative features were selected as the 8 RNA genes with the largest β row L2 norm across the 221 gene-target perturbations.

Gene labels were colored by bulk MAGeCK classification. Genes classified as HLA activators, defined by top-100 enrichment in HLA-Low versus Input, were colored blue. Genes classified as HLA repressors, defined by top-100 enrichment in HLA-High versus Input, were colored red. Genes not meeting either criterion were colored black.

Edges between co-functional modules and gene programs summarize aggregate directional effects. For each co-functional module *M* and gene program *P*, the mean β value was computed over the submatrix whose rows belong to *P* and whose columns belong to *M*.

### Mechanistic interpretation and agentic-assisted regulatory modeling

To integrate phenotypic enrichment, single-cell perturbation effects and transcriptomic response programs into a mechanistic model, Google DeepMind’s Co-Scientist was used as an AI-assisted interpretation framework. Co-Scientist was provided with the seven co-functional modules, the top 25 representative genes from each of the nine gene programs, and a table of the top 100 HLA activator and repressor hits from the MAGeCK bulk CRISPR screen. Notably, 49 of these activators and 17 of these repressors were included among the 221 genes retained in the single-cell regularized model. For each MAGeCK hit, the input included its enrichment ranking, log-fold change, and its association with the co-functional modules (co-functional module ID) defined in the regulatory model. Co-Scientist was asked to annotate the co-functional modules and gene programs, cross-reference existing literature and prioritize candidate mechanisms linking perturbation modules to HLA-I surface expression (**Supplementary Material**).

Ten significant MAGeCK hits (FDR < 0.05) that were not included among the 221 retained gene-target perturbations in the regulatory model were treated as high-confidence orphan hits. These genes were not used to define co-functional modules or gene programs, but were provided as input to Co-Scientist to be considered during mechanistic interpretation and mapped to regulatory axes when supported by literature-derived links.

Expert manual review was used to consolidate the Co-Scientist’s output, and extract overarching biological themes including IFNγ signaling, transcriptional regulation, biosynthesis, proteostasis, ER quality control, Golgi trafficking, retromer-dependent recycling and surface HLA-I regulation. For clarity, Co-Scientist generated these specific mechanistic groupings and used these exact keywords (e.g., ‘Proteostasis Axis’, ‘Golgi/Trafficking Axis’, ‘ER quality control’) in its narrative output. Human scientists extracted and formatted these words into the final clean list of labels.

### Regulatory-network visualization and spatial pathway mapping

To visualize the inferred regulatory architecture of HLA-I control, a hierarchical regulatory network was constructed using a custom Python workflow based on Plotly. The Sankey diagram summarized the inferred flow from genetic perturbation modules to RNA gene programs and terminal HLA-I surface-expression phenotypes. Network nodes were defined as the seven co-functional modules, the nine gene programs and terminal HLA-Low or HLA-High phenotype states.

Edges from co-functional modules to gene programs were weighted by the mean ElasticNet coefficient β across all target-feature pairs connecting a given co-functional module to a given gene program. Module-to-phenotype direction was assigned according to bulk enrichment behavior. Perturbations whose knockout decreased surface HLA-I were assigned to the activation axis, indicating that the targeted genes normally support HLA-I expression. Perturbations whose knockout increased surface HLA-I were assigned to the repression or buffering axis, indicating that the targeted genes normally restrain HLA-I surface abundance.

To translate this network into a biologically interpretable cellular model, co-functional modules and gene programs were mapped onto a spatial schematic of MHC-I biogenesis. Each co-functional module was assigned to candidate cellular processes and subcellular locations based on the known functions and consensus localization of its constituent gene products, using Human Protein Atlas annotations and literature-supported functional evidence. Gene programs were mapped according to their representative marker genes and pathway labels.

Using BioRender, the Co-Scientist-derived regulatory model was integrated into this spatial framework. Orphan hits from the MAGeCK screen were overlaid at predicted functional sites when supported by literature-derived mechanistic links.

## CODE AVAILABILITY

All analysis code is available at https://github.com/Genentech/PerturbME.

## DATA AVAILABILITY

Analysis-ready single-cell objects, the perturbation-effect model and its module and gene-program assignments are available at figshare (https://doi.org/10.6084/m9.figshare.33264225). Raw and processed sequencing data have been deposited in the NCBI Gene Expression Omnibus and will be available before publication. Source data are provided with this paper.

## ACKNOWLEDGEMENTS

We thank Leslie Gaffney for help with figure design. lentiCas9-Blast was a gift from Feng Zhang (Addgene 52962).

## COMPETING INTERESTS

HW, JG, AS, ES, OR-R, KG-S, PIT and AR are employees of Genentech (a member of the Roche Group) or Roche, and AS, ES, OR-R, KG-S, PIT and AR have equity in Roche. ZX was a paid intern in Genentech during the study. TB and VN are employees of Google/Alphabet and may own Alphabet stock as part of the standard compensation package. A.R., O.R.R., K.G.S., and P.I.T. are inventors on multiple patents filed by or issued to the Broad Institute related to single cell genomics and Perturb-Seq including a patent application related to Perturb-ME.

---

**Extended Data Table 1.**
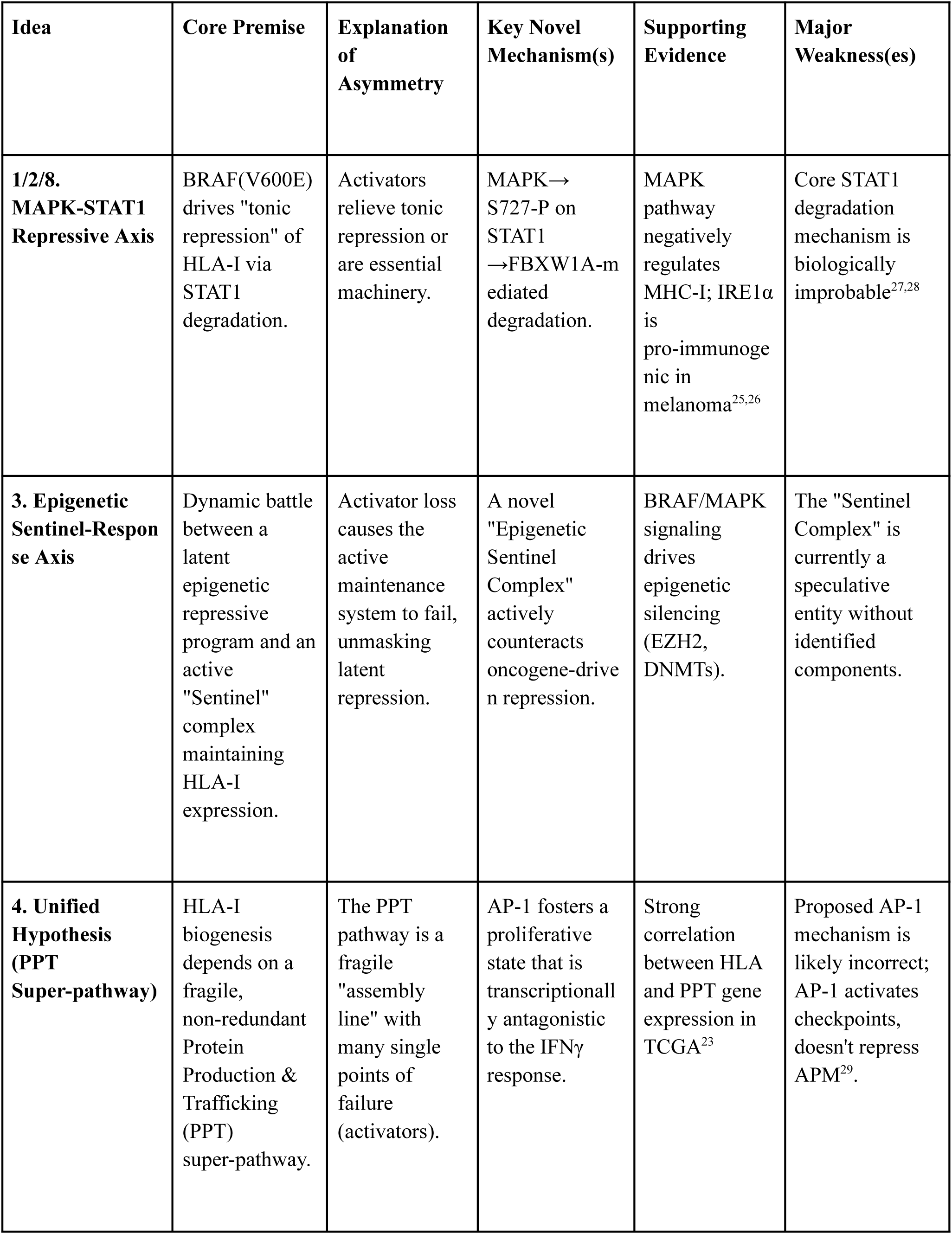
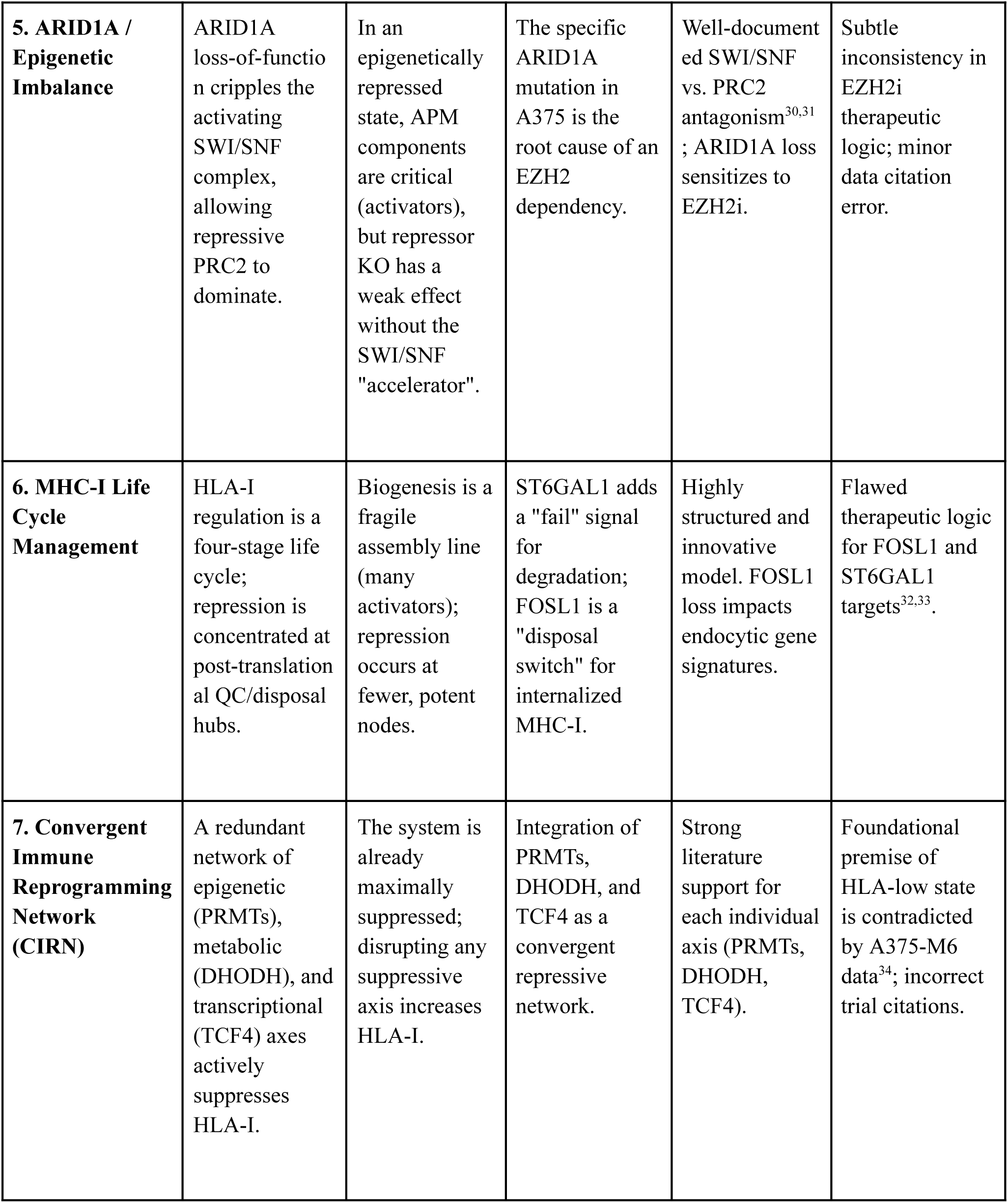
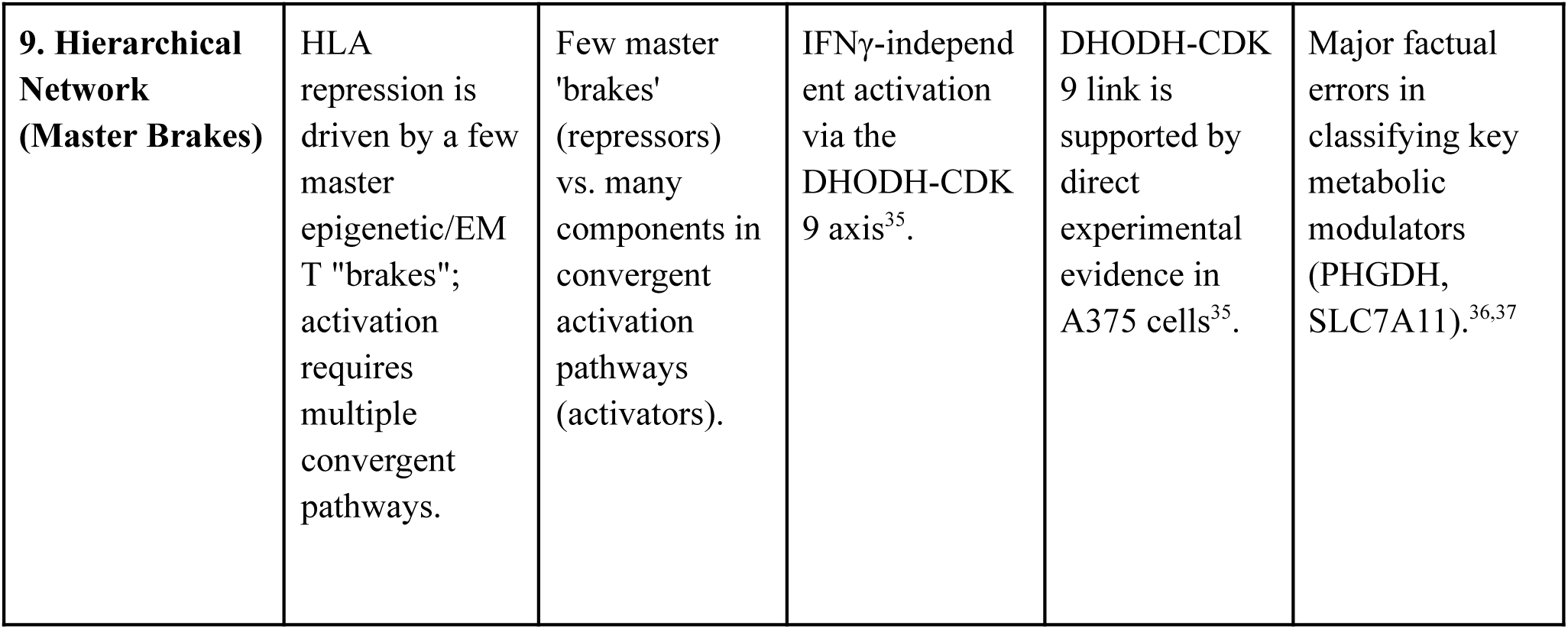
Co-scientist Hypothesis Tournament and Deep Verification.

| **Idea** | **Core Premise** | **Explanation of Asymmetry** | **Key Novel Mechanism(s)** | **Supporting Evidence** | **Major Weakness(es)** |
| --- | --- | --- | --- | --- | --- |
| **1/2/8. MAPK-STAT1 Repressive Axis** | BRAF(V600E) drives "tonic repression" of HLA-I via STAT1 degradation. | Activators relieve tonic repression or are essential machinery. | MAPK→S727-P on STAT1 →FBXW1A-mediated degradation. | MAPK pathway negatively regulates MHC-I; IRE1α is pro-immunogenic in melanoma<sup>25,26</sup> | Core STAT1 degradation mechanism is biologically improbable<sup>27,28</sup> |
| **3. Epigenetic Sentinel-Response Axis** | Dynamic battle between a latent epigenetic repressive program and an active "Sentinel" complex maintaining HLA-I expression. | Activator loss causes the active maintenance system to fail, unmasking latent repression. | A novel "Epigenetic Sentinel Complex" actively counteracts oncogene-driven repression. | BRAF/MAPK signaling drives epigenetic silencing (EZH2, DNMTs). | The "Sentinel Complex" is currently a speculative entity without identified components. |
| **4. Unified Hypothesis (PPT Super-pathway)** | HLA-I biogenesis depends on a fragile, non-redundant Protein Production & Trafficking (PPT) super-pathway. | The PPT pathway is a fragile "assembly line" with many single points of failure (activators). | AP-1 fosters a proliferative state that is transcriptionally antagonistic to the IFNγ response. | Strong correlation between HLA and PPT gene expression in TCGA<sup>23</sup> | Proposed AP-1 mechanism is likely incorrect; AP-1 activates checkpoints, doesn't repress APM<sup>29</sup>. |
| **5. ARID1A / Epigenetic Imbalance** | ARID1A loss-of-function cripples the activating SWI/SNF complex, allowing repressive PRC2 to dominate. | In an epigenetically repressed state, APM components are critical (activators), but repressor KO has a weak effect without the SWI/SNF "accelerator". | The specific ARID1A mutation in A375 is the root cause of an EZH2 dependency. | Well-documented SWI/SNF vs. PRC2 antagonism<sup>30,31</sup>; ARID1A loss sensitizes to EZH2i. | Subtle inconsistency in EZH2i therapeutic logic; minor data citation error. |
| **6. MHC-I Life Cycle Management** | HLA-I regulation is a four-stage life cycle; repression is concentrated at post-translational QC/disposal hubs. | Biogenesis is a fragile assembly line (many activators); repression occurs at fewer, potent nodes. | ST6GAL1 adds a "fail" signal for degradation; FOSL1 is a "disposal switch" for internalized MHC-I. | Highly structured and innovative model. FOSL1 loss impacts endocytic gene signatures. | Flawed therapeutic logic for FOSL1 and ST6GAL1 targets<sup>32,33</sup>. |
| **7. Convergent Immune Reprogramming Network (CIRN)** | A redundant network of epigenetic (PRMTs), metabolic (DHODH), and transcriptional (TCF4) axes actively suppresses HLA-I. | The system is already maximally suppressed; disrupting any suppressive axis increases HLA-I. | Integration of PRMTs, DHODH, and TCF4 as a convergent repressive network. | Strong literature support for each individual axis (PRMTs, DHODH, TCF4). | Foundational premise of HLA-low state is contradicted by A375-M6 data<sup>34</sup>; incorrect trial citations. |
| **9. Hierarchical Network (Master Brakes)** | HLA repression is driven by a few master epigenetic/EM T "brakes"; activation requires multiple convergent pathways. | Few master 'brakes' (repressors) vs. many components in convergent activation pathways (activators). | IFNγ-independent activation via the DHODH-CDK 9 axis<sup>35</sup>. | DHODH-CDK 9 link is supported by direct experimental evidence in A375 cells<sup>35</sup>. | Major factual errors in classifying key metabolic modulators (PHGDH, SLC7A11).<sup>36,37</sup> |

## SUPPLEMENTARY FILES

Supplementary File 1 - Co-Scientist prompt

Supplementary File 2 - Co-Scientist response

Supplementary File 3 - MAGeCK bulk screen analysis (zipped folder)

Supplementary Table 1 - CITE-Seq antibody panel

Supplementary Table 2 - CRISPR guide library

Supplementary Table 3 - Bulk enrichment *vs*. Perturb-ME concordance

Supplementary Table 4 - Co-functional module and gene-program gene lists

Supplementary Table 5 - Guide-seq primer

## Notes

https://doi.org/10.6084/m9.figshare.33264225

https://github.com/Genentech/PerturbME

