## Supplementary material for "Perturb-ME: Scalable mechanism discovery from phenotype-enriched genome-wide screens": Supplementary File 1 - CoScientist Prompt.docx

### **Revised Prompt for Co-Sciencist: Mechanistic Interpretation of the HLA Regulatory Network**

**Written by**: Jiacheng Gu

**Last updated**: Feb. 13, 2026

**Goal:** Provide a comprehensive biological annotation of the regulatory network controlling MHC Class I (HLA) in A375 melanoma cells. The Co-scientist must interpret the "global landscape" by mapping high-impact hits from bulk CRISPR screens to the functional "Output Programs" defined by single-cell CITE-seq.

#### **I. Biological Context & Experimental System**

1. **System**: A375 human melanoma cells (BRAF V600E) treated with IFNγ (2 ng/mL, 24 hours) to induce baseline HLA expression
2. **Perturbation**: Genome-wide CRISPR knockout library (19,767 genes, 3 sgRNAs per gene, plus 1,000 non-targeting and 1,000 intergenic controls).
3. **Enrichment Strategy**: Cells were FACS-sorted into **HLA_Low (Bottom 5%)** and **HLA_High (Top 5%)** populations based on HLA-A,B,C surface expression prior to single-cell profiling. This design ensures that the data is enriched for perturbations with a high phenotypic impact on HLA surface density.
4. **Single-Cell Readout**
   - RNA: ~18,670 genes per cell
   - Protein: 13 surface markers via CITE-seq antibodies:
     - HLA presentation: HLA-A,B,C (CITE-HLA_A), HLA-E, HLA-DR
     - Immune signaling: CD274 (PD-L1), CD119 (IFNγR), CD47, CD58
     - Other: CD29, CD44, CD49f, CD59, CD61, CD107a

#### **II. Initial data generation**

1. **Regulation Matrix (Primary Data for Interpretation):** To quantify the specific effect of each gene knockout on the transcriptomic and proteomic features, we trained an **ElasticNet Linear Model**. The resulting **Beta Matrix** contains the ElasticNet regression coefficients (β) representing the weight of every gene-to-feature relationship.
   - Positive β: Knocking out this target INCREASES feature expression
   - Negative β: Knocking out this target DECREASES feature expression
2. **Clustering:** We applied dimensionality reduction to the Beta Matrix to group genes into two types of modules:
   - **Target Clusters:** Groups of **perturbations (knockouts)** by similarity of their regulatory profiles (β correlation), i.e. groups of perturbations that cause similar cellular responses. **7 target modules** were identified.
   - **Feature Clusters:** Groups of **RNAs/proteins (readouts)** by co-regulation pattern, i.e. RNAs/proteins that respond coordinately to the same perturbations.**9 feature clusters** were identified.
3. **Bulk gRNA Enrichment (MAGeCK)**: perturbed cell population was sorted via FACS into distinct populations based on surface HLA expression. Genomic DNA was extracted and sgRNA abundance was quantified for:
   - **HLA_High (Top 5%)**: Enriched for cells where the knockout increased HLA expression.
   - **HLA_Low (Bottom 5%):** Enriched for cells where the knockout decreased HLA expression.
   - **Input**: The unsorted control population representing the starting library distribution.

The MAGeCK algorithm compares the sorted gates to the Input pool to calculate two key parameters for each gene:

- **RRA Score** (Robust Rank Aggregation): A significance metric (similar to a p-value). A lower RRA score indicates that multiple independent sgRNAs for a gene are consistently enriched, suggesting a robust biological effect.
- **Log-Fold Change** (LFC): The magnitude of enrichment. A positive LFC in a specific gate indicates the gene knockout is highly prevalent in that population.

The data is divided into two primary comparisons that define the role of the gene in the HLA pathway:

- **High vs. Input (The Repressors)**: Genes with high significance in this comparison are HLA Repressors. Their normal function is to keep HLA levels in check; when knocked out, HLA levels rise.
- **Low vs. Input (The Activators)**: Genes with high significance in this comparison are HLA Activators. These are essential components of the MHC I machinery; when knocked out, the cell loses its ability to express HLA.

#### **III. Input for Co-Scientist (Processed for Interpretability)**

Because the raw data is high-dimensional and contains technical noise, we applied three processing steps to make it readable for the Co-scientist:

**Step 1: Selecting the Top 100 MAGeCK Hits**

We filtered the MAGeCK data to select the **Top 100 Repressors** and **Top 100 Activators** based on statistical significance (RRA score). This enriches genes with the strongest biological impact on HLA expression level.

**Step 2: Summarizing Features by top 25 representative genes**

We used the beta matrix to calculate the **L2 Norm** for every gene. We then selected the **Top 25 most responsive genes** for each feature cluster. These represent the most robust "sensors" of the cellular program.

**Step 3: Mapping enriched gRNA to target modules**

We cross-referenced the Top 100 MAGeCK high or low hits with the list of target clusters.

- We highlighted hits that meet the **> 17 cells per target** threshold and are part of a **target cluster**. These are high-confidence hits.
- The remaining MAGeCK hits will be **Discovery Frontiers.** They are genes that had massive phenotypic impact in the bulk gRNA enrichment but lacked sufficient cell coverage to be mapped to a target cluster.

##### **Input Data 1: List of 224 genes in 7 Target Clusters grouped by similarity of regulatory profiles**

**Cluster 0 (48 genes)**: ARMCX2, B2M, B3GAT1, BRMS1, C12orf50, C1GALT1C1, CCDC153, CMAS, DECR1, DEGS2, DMRTC2, E2F2, EED, FOSL1, GPR174, HAS1, HIGH_CTL, HNRNPM, HTT, KMT5C, LOC107986211, MAGEB6, MGAT1, MYEF2, NTAN1, OR5J2, PNMA5, RAD51AP1, RBM10, RBSN, RRAGB, SETD1B, SETD2, SH2D5, SLAIN1, SMIM10L2A, SPTLC3, SUSD6, TADA1, TAF5L, TAP1, TAPBP, TM2D3, TMEM258, VPS29, VPS35, WNT10B, WWP2

**Cluster 1 (23 genes)**: AATF, C7orf26, COPS3, COPS5, COPS8, DBR1, DIMT1, DPAGT1, DUSP4, EIF4G1, ELP5, INTS1, MCM3AP, MED20, NUDT21, PEA15, POLR3B, POLR3F, PSMG4, SNRNP40, SOD1, URM1, ZNF236

**Cluster 2 (53 genes)**: ALG1, AMH, ANKS1A, ANTKMT, ARSK, ASB7, ASCC3, BEST3, C1QL3, C5orf30, CCNJL, CFB, CIITA, CRYGB, CYB5R4, CYTL1, DDHD2, DDX39B, DET1, FKBP5, GPR107, INHA, IRF1, ISOC1, ITGA6, JUP, KRTAP12-3, L3MBTL3, LOC105379752, MED16, MFSD4B, MS4A7, NEK6, NLRC5, NYAP2, OR2J2, PLPBP, PMPCB, PRR5, RAB44, RFX5, RFXANK, RFXAP, RSPH4A, SMAP1, SOBP, SPPL3, SRSF6, TBC1D25, TMEM207, TMEM242, TREML2, TRIP12

**Cluster 3 (27 genes)**: ALG2, BRF1, CDC123, ELAC2, ELP1, GRPEL1, GTF3C3, HARS1, IARS1, LAGE3, LOW_CTL, NUS1, OSGEP, POLR3E, POLR3H, PREB, RABGGTA, RPP21, RTCB, SPCS2, TBP, TRMT5, VARS, YARS1, YRDC, ZBTB8OS, ZPR1

**Cluster 4 (44 genes)**: ARFRP1, BOK, C7orf57, CALR, CHST13, COL6A3, CREBBP, ENPP1, EXOSC10, GADD45B, GALNT1, GSK3B, HLA-B, HLA-C, INSM1, JAG2, KLHL30, KRTAP12-4, LCE5A, LEFTY2, LOC112267886, MDK, MPIG6B, MRPS18B, NANOGNB, NANOS2, NCSTN, OR10H2, OR5M3, OR5T2, OTUD5, PASD1, RGS9BP, SATL1, SCARA5, SLC35A1, SLC35A2, SLC39A12, SMIM19, TAP2, TNNT1, TSPAN4, USP41, ZNF784

**Cluster 5 (13 genes)**: GTF3C4, IFNGR1, IFNGR2, JAK1, JAK2, SRP14, SRP19, SRP54, SRP68, SRP72, SRP9, SRPRA, STAT1

**Cluster 6 (16 genes**): ARF4, CAMLG, COG1, COG3, COG4, COG8, G1, GET1, GET3, NMT1, SLC39A7, SPCS3, SYS1, TMED10, UNC50, USO1

##### **Input Data 2: Top 100 HLA Activators Identified via Bulk MAGeCK Enrichment (HLA-Low vs. Input)**

| **Gene Name** | **Enrichment Ranking** | **Log Fold-Change (LFC)** | **In Target Cluster?** | **Target Cluster ID** |
| --- | --- | --- | --- | --- |
| TAP2 | 1 | 5.2834 | TRUE | 4 |
| TAPBP | 2 | 4.6293 | TRUE | 0 |
| TAP1 | 3 | 4.7484 | TRUE | 0 |
| IFNGR2 | 4 | 4.5855 | TRUE | 5 |
| JAK2 | 5 | 4.5829 | TRUE | 5 |
| STAT1 | 6 | 4.4887 | TRUE | 5 |
| SRPRA | 7 | 4.1856 | TRUE | 5 |
| JAK1 | 8 | 3.9842 | TRUE | 5 |
| RFXAP | 9 | 3.7881 | TRUE | 2 |
| TAF2 | 10 | 4.7231 | FALSE |  |
| IRF1 | 11 | 4.6202 | TRUE | 2 |
| ALG11 | 12 | 3.2645 | FALSE |  |
| B2M | 13 | 3.3346 | TRUE | 0 |
| DHDDS | 14 | 3.2134 | FALSE |  |
| SYS1 | 15 | 4.1546 | TRUE | 6 |
| RABGGTA | 16 | 3.6909 | TRUE | 3 |
| SPPL3 | 17 | 3.8888 | TRUE | 2 |
| CHAF1B | 18 | 3.2671 | FALSE |  |
| POLR3H | 19 | 3.2725 | TRUE | 3 |
| SLC39A7 | 20 | 3.3315 | TRUE | 6 |
| COPS8 | 21 | 2.9145 | TRUE | 1 |
| SRP14 | 22 | 4.7233 | TRUE | 5 |
| ARF4 | 23 | 4.7713 | TRUE | 6 |
| IFNGR1 | 24 | 4.6182 | TRUE | 5 |
| HARS1 | 25 | 3.8562 | TRUE | 3 |
| ELAC2 | 26 | 3.2403 | TRUE | 3 |
| NLRC5 | 27 | 3.1276 | TRUE | 2 |
| NUS1 | 28 | 4.2486 | TRUE | 3 |
| MPHOSPH10 | 29 | 4.4538 | FALSE |  |
| RPP21 | 30 | 4.7768 | TRUE | 3 |
| RFXANK | 31 | 2.8773 | TRUE | 2 |
| FAM210A | 32 | 2.7166 | FALSE |  |
| GTF3C3 | 33 | 3.1913 | TRUE | 3 |
| SLC20A2 | 34 | 2.658 | FALSE |  |
| SRP54 | 35 | 4.5191 | TRUE | 5 |
| ZBTB8OS | 36 | 3.0251 | TRUE | 3 |
| EIF2B3 | 37 | 3.4665 | FALSE |  |
| HMGCS1 | 38 | -6.6452 | FALSE |  |
| AFG3L2 | 39 | 3.3859 | FALSE |  |
| SRP9 | 40 | 2.8399 | TRUE | 5 |
| GMPPB | 41 | 3.7416 | FALSE |  |
| FARSB | 42 | 2.9606 | FALSE |  |
| CARS1 | 43 | 4.5859 | FALSE |  |
| CALR | 44 | 3.2658 | TRUE | 4 |
| ELP3 | 45 | 4.9871 | FALSE |  |
| DPH3 | 46 | 3.4241 | FALSE |  |
| LLGL2 | 47 | 0.91133 | FALSE |  |
| OTUD5 | 48 | 2.1905 | TRUE | 4 |
| CDC123 | 49 | 2.6383 | TRUE | 3 |
| SGK1 | 50 | 2.627 | FALSE |  |
| TEPSIN | 51 | 3.0108 | FALSE |  |
| OSGEP | 52 | 2.9127 | TRUE | 3 |
| RRP36 | 53 | 3.5582 | FALSE |  |
| TRMT5 | 54 | 3.1465 | TRUE | 3 |
| WDR92 | 55 | 3.2188 | FALSE |  |
| FBXO36 | 56 | 2.5342 | FALSE |  |
| KRTAP19-1 | 57 | 2.4293 | FALSE |  |
| PYROXD1 | 58 | 3.5857 | FALSE |  |
| MED20 | 59 | 2.8578 | TRUE | 1 |
| COPS6 | 60 | 3.8344 | FALSE |  |
| XRCC6 | 61 | 3.8688 | FALSE |  |
| RFX5 | 62 | 3.2898 | TRUE | 2 |
| SSB | 63 | 2.6018 | FALSE |  |
| C7orf26 | 64 | 2.392 | TRUE | 1 |
| FAM120AOS | 65 | 0.39162 | FALSE |  |
| NRBP1 | 66 | 3.2976 | FALSE |  |
| NOP10 | 67 | 2.8239 | FALSE |  |
| C1orf131 | 68 | 3.1077 | FALSE |  |
| ZNHIT6 | 69 | 2.7405 | FALSE |  |
| USP45 | 70 | 2.5555 | FALSE |  |
| SRP68 | 71 | 3.1178 | TRUE | 5 |
| RNMT | 72 | 2.7111 | FALSE |  |
| CREBBP | 73 | 2.3617 | TRUE | 4 |
| THOC1 | 74 | 3.4386 | FALSE |  |
| SOD1 | 75 | 2.4676 | TRUE | 1 |
| CHSY3 | 76 | 2.1414 | FALSE |  |
| COPS5 | 77 | 2.7519 | TRUE | 1 |
| UTP25 | 78 | 3.1294 | FALSE |  |
| ELP5 | 79 | 3.3007 | TRUE | 1 |
| RPP38 | 80 | 3.0233 | FALSE |  |
| GBP1 | 81 | 1.5256 | FALSE |  |
| GSTA3 | 82 | 2.9985 | FALSE |  |
| CMTR1 | 83 | 2.9771 | FALSE |  |
| NUDT21 | 84 | 2.5599 | TRUE | 1 |
| VARS | 85 | 3.3724 | TRUE | 3 |
| LOC102724159 | 86 | 3.521 | FALSE |  |
| ABT1 | 87 | 3.1468 | FALSE |  |
| PPIE | 88 | 2.7952 | FALSE |  |
| INTS7 | 89 | 3.7779 | FALSE |  |
| PREB | 90 | 3.6215 | TRUE | 3 |
| AGPAT1 | 91 | 2.4486 | FALSE |  |
| AP5S1 | 92 | 1.1556 | FALSE |  |
| CEBPZ | 93 | 4.5236 | FALSE |  |
| TBP | 94 | 2.9776 | TRUE | 3 |
| DDX39B | 95 | 2.559 | TRUE | 2 |
| GJD3 | 96 | 2.0338 | FALSE |  |
| RRP7A | 97 | 2.4364 | FALSE |  |
| ALG2 | 98 | 3.3482 | TRUE | 3 |
| HYOU1 | 99 | 3.2649 | FALSE |  |
| GUK1 | 100 | 3.2221 | FALSE |  |

##### **Input Data 3: Top 100 HLA Repressors Identified via Bulk MAGeCK Enrichment (HLA-High vs. Input)**

| **Gene Name** | **Enrichment Ranking** | **Log Fold-Change (LFC)** | **In Target Cluster?** | **Target Cluster ID** |
| --- | --- | --- | --- | --- |
| VPS35 | 1 | 4.7004 | TRUE | 0 |
| SLC35A1 | 2 | 4.0095 | TRUE | 4 |
| COG3 | 3 | 4.432 | TRUE | 6 |
| F8A2 | 4 | 3.7 | FALSE |  |
| COG1 | 5 | 4.4267 | TRUE | 6 |
| GATA2 | 6 | 3.1482 | FALSE |  |
| C1GALT1 | 7 | 3.94 | FALSE |  |
| FOSL1 | 8 | 4.4182 | TRUE | 0 |
| MAPK3 | 9 | 3.6119 | FALSE |  |
| SNRNP200 | 10 | -5.3179 | FALSE |  |
| JUND | 11 | 3.718 | FALSE |  |
| MARK2 | 12 | 4.0618 | FALSE |  |
| TAZ | 13 | 3.2739 | FALSE |  |
| PRR3 | 14 | 1.8708 | FALSE |  |
| RBM10 | 15 | 3.6966 | TRUE | 0 |
| SCIMP | 16 | 4.113 | FALSE |  |
| COG8 | 17 | 4.0285 | TRUE | 6 |
| NUTF2 | 18 | 3.0539 | FALSE |  |
| SOAT1 | 19 | 3.9534 | FALSE |  |
| BBOF1 | 20 | 3.9193 | FALSE |  |
| TMEM101 | 21 | 3.8604 | FALSE |  |
| LRG1 | 22 | 3.1111 | FALSE |  |
| PKNOX2 | 23 | 3.1214 | FALSE |  |
| GALNT1 | 24 | 3.6687 | TRUE | 4 |
| ARSG | 25 | 3.3583 | FALSE |  |
| CMAS | 26 | 3.7937 | TRUE | 0 |
| COMMD3-BMI1 | 27 | 3.1119 | FALSE |  |
| WRB-SH3BGR | 28 | 3.2774 | FALSE |  |
| SETD2 | 29 | 3.5975 | TRUE | 0 |
| STAG2 | 30 | 3.3797 | FALSE |  |
| TAF1C | 31 | 3.993 | FALSE |  |
| LSM7 | 32 | 2.0347 | FALSE |  |
| NISCH | 33 | 3.1642 | FALSE |  |
| KIAA1328 | 34 | 2.9632 | FALSE |  |
| VSTM5 | 35 | 2.4026 | FALSE |  |
| SMIM3 | 36 | 4.118 | FALSE |  |
| CLASP1 | 37 | 3.1635 | FALSE |  |
| MYH9 | 38 | 3.5201 | FALSE |  |
| HSF4 | 39 | 4.254 | FALSE |  |
| TADA2B | 40 | 2.8719 | FALSE |  |
| ZCCHC14 | 41 | 3.2515 | FALSE |  |
| BIRC2 | 42 | 4.1 | FALSE |  |
| NDUFB1 | 43 | 3.3988 | FALSE |  |
| TGM4 | 44 | 3.3313 | FALSE |  |
| PATE4 | 45 | 3.4738 | FALSE |  |
| CAMLG | 46 | 3.8018 | TRUE | 6 |
| LUM | 47 | 3.588 | FALSE |  |
| PAPOLA | 48 | 4.0077 | FALSE |  |
| GET3 | 49 | 3.8238 | TRUE | 6 |
| RAB28 | 50 | 3.004 | FALSE |  |
| C5orf46 | 51 | 0.31935 | FALSE |  |
| CEP41 | 52 | 3.8738 | FALSE |  |
| LRMP | 53 | 3.398 | FALSE |  |
| RPL18A | 54 | 3.6382 | FALSE |  |
| MAPK1 | 55 | 3.4454 | FALSE |  |
| KISS1 | 56 | 3.5665 | FALSE |  |
| C16orf82 | 57 | 3.0553 | FALSE |  |
| BICDL1 | 58 | 3.0841 | FALSE |  |
| SLC35A2 | 59 | 3.2665 | TRUE | 4 |
| MYH13 | 60 | 3.6289 | FALSE |  |
| SNCG | 61 | 1.6515 | FALSE |  |
| LOC105370092 | 62 | 3.3958 | FALSE |  |
| GLT8D1 | 63 | 3.1225 | FALSE |  |
| ZNF684 | 64 | 1.5271 | FALSE |  |
| KIF5C | 65 | 3.6798 | FALSE |  |
| DACH2 | 66 | 2.9419 | FALSE |  |
| ZNF595 | 67 | 3.4644 | FALSE |  |
| MYO1H | 68 | 2.9749 | FALSE |  |
| FUCA2 | 69 | 3.3117 | FALSE |  |
| MYC | 70 | 3.7224 | FALSE |  |
| KCTD7 | 71 | 2.9416 | FALSE |  |
| PIGY | 72 | 3.2849 | FALSE |  |
| HAL | 73 | 2.9783 | FALSE |  |
| NME5 | 74 | 3.0799 | FALSE |  |
| TUBGCP6 | 75 | 3.7849 | FALSE |  |
| CSRP3 | 76 | 1.2322 | FALSE |  |
| C2orf83 | 77 | 3.1211 | FALSE |  |
| HDAC6 | 78 | 2.3551 | FALSE |  |
| ZNF195 | 79 | 3.4028 | FALSE |  |
| TSSK4 | 80 | 3.8332 | FALSE |  |
| VPS29 | 81 | 3.2877 | TRUE | 0 |
| LAX1 | 82 | 3.724 | FALSE |  |
| PDXDC1 | 83 | 3.5047 | FALSE |  |
| MUC19 | 84 | 3.3496 | FALSE |  |
| SPACA5 | 85 | 2.4686 | FALSE |  |
| WDR13 | 86 | 3.5875 | FALSE |  |
| OR51A7 | 87 | 2.728 | FALSE |  |
| TRIM42 | 88 | 3.4865 | FALSE |  |
| C14orf180 | 89 | 2.8541 | FALSE |  |
| LSM3 | 90 | 0.56614 | FALSE |  |
| CAPN1 | 91 | 3.1895 | FALSE |  |
| ROS1 | 92 | 3.6076 | FALSE |  |
| STXBP4 | 93 | 2.1443 | FALSE |  |
| NME7 | 94 | 3.0524 | FALSE |  |
| LOC105376525 | 95 | 0.69811 | FALSE |  |
| C1GALT1C1 | 96 | 3.1019 | TRUE | 0 |
| MGAT1 | 97 | 3.3798 | TRUE | 0 |
| WWP2 | 98 | 3.0414 | TRUE | 0 |
| PHKG1 | 99 | 1.5208 | FALSE |  |
| DDRGK1 | 100 | 3.163 | FALSE |  |

##### **Input Data 4: top 25 representative genes of 9 feature clusters**

**Cluster 0:** S100B, FXYD3, NME4, HAPLN1, MARCKSL1, SOX5, SLC26A2, CTSZ, MDK, GSN, CYBA, S100A13, CYB5A, VIM, CALM2, GNG12, LGALS1, EMP3, CTNNB1, COL6A3, TIMP1, CD63, S100A6, S100A10, MGST1.

**Cluster 1:** MANF, SDF2L1, ARF4, RABAC1, HSPA5, YIF1A, LMAN1, KDELR2, CRELD2, ERLEC1, P4HB, CALU, SEC11C, DNAJB11, SRPRB, PDIA6, HSP90B1, CANX, RPN1, SEC61A1, DDX39B, EIF2B2, DNAJC3, TRAM1, GANAB.

**Cluster 2:** IL24, GPRC5A, SLC3A2, DDIT3, MTHFD2, PSAT1, PHGDH, CD55, HERPUD1, DDIT4, ASNS, CEBPB, TRIB3, TENT5A, ATF3, SLC7A11, PCK2, CARS1, SLC1A5, SARS1, GPT2, STC2, ATF4, GADD45A, TFF3.

**Cluster 3:** ID3, TGFA, STK17A, SNHG25, CAVIN1, NFIC, MSN, SLC5A3, IGFBP6, ALDH1A3, AHNAK, COL1A1, THBS1, EMP1, SERPINE1, ANXA2, ITGA2, FOS, JUN, CCN1, JUNB, MYC, EGR1, IER3, KLF4.

**Cluster 4:** FST, ID1, S100A2, PYGB, MOK, RAD23A, PLP2, EIF2S3, MCL1, TMEM230, GADD45B, ATF5, PPP1R15A, PMAIP1, BBC3, BAX, CDKN1A, MDM2, FAS, BTG2, SESN2, PLK2, GDF15, CCNG1, PHLDA3.

**Cluster 5:** AKAP12, NNMT, CAV1, MT-ND6, HSPB1, PRSS23, TPM1, TIMP3, BAX, SNRNP70, GAPVD1, FLNA, MYH9, ACTB, ACTG1, PFN1, CFL1, ARPC2, CAP1, DSTN, VCL, TLN1, IQGAP1, ANXA1, ANXA6.

**Cluster 6:**MRPS6, RPL22L1, MFSD12, TMEM179B, TMSB4X, BNIP3, PHLDA2, TIMM8B, HIST1H4C, ERCC1, EIF4EBP1, RPS6, RPL11, EEF1A1, EEF2, RPS3, RPL7, RPS18, RPLP1, RPS2, RPL13A, RPS4X, RPL19, RPS13, RPL21.

**Cluster 7:** BST2, GBP1, HLA-C, STAT1, HLA-DMA, IRF1, VAMP5, HLA-DRB5, UBE2L6, PSMB9, TAP1, HLA-B, B2M, PSME2, HLA-A, TAPBP, NLRC5, PSMB8, IFI30, CD74, PSME1, HLA-DPA1, HLA-DPB1, HLA-DQB1, HLA-DRA.

**Cluster 8:** PMEPA1, PPDPF, FLNA, LRPPRC, CSKMT, PHLDA3, DUSP4, ATP1B3, CAPG, BRIX1, WDR12, PES1, BOP1, NOLC1, EBNA1BP2, BYSL, GNL3, UTP14A, WDR75, PWP2, RRP12, NOB1, NAT10, DDX21, RRP1.

#### **IV. Guideline for the Co-Scientist**

**Overall objective:** Establish a comprehensive end-to-end hypothesis of HLA regulation by integrating global phenotypic drivers with high-resolution transcriptional programs.

##### **1. Functional Annotation of Target Clusters**

First, perform a high-level summary of the 7 Target Clusters provided above. Identify the biological "pathway identity" of each cluster (e.g., *Is Cluster 6 representing the COG/Golgi machinery? Is Cluster 5 the JAK/STAT signaling axis?*). Note that Cluster 0 contains the "High Control" and Cluster 3 contains the "Low Control," which serve as phenotypic landmarks.

##### **2. Solving the "Discovery Frontier"**

Analyze the **Top 100 HLA Repressors (High vs. Input)** and **Top 100 HLA Activators (Low vs. Input)**.

- Identify high-ranking genes that are **not** in the 224 mapped targets (Input Data 1).
- Can these novel hits be conceptually mapped to the existing 7 programs? (e.g., Is a novel hit likely a Golgi transport factor?)
- Are there entirely new regulatory programs emerging (e.g., RNA splicing or metabolic dependencies)?

##### **3. Connecting Perturbations to Feature Programs**

Examine the **9 Feature Modules** (Output Programs).

- Summarize the biological theme of each module based on the Top 25 Priority Features (e.g., Cluster 7 = Antigen Presentation).
- Establish the links: Explain how specific **Target Clusters** (Inputs) selectively modulate these **Feature Programs** (Outputs).
- Specifically, address how these shifts lead to the final change in surface HLA expression (e.g., transcriptional activation vs. post-translational stability).

##### **4. Synthesis: The End-to-End Hypothesis**

Build a cohesive narrative for the major regulatory axes identified.

- **Logic:** Target Program (e.g., Retrograde Transport) → Cellular Response Program (e.g., Integrated Stress) → Final Phenotype (HLA-High/Low).

##### **5. Extra Credit: Therapeutic Targeting & Evidence**

Identify specific nodes in the identified regulatory network that could serve as therapeutic targets for restoring immunogenicity in "cold" tumors. Provide **PMID citations** and reference public datasets (including melanoma-specific or non-melanoma context) to support your therapeutic hypotheses.
